# Pancreatic islet cell stress induced by insulin-degrading enzyme deficiency promotes islet regeneration and protection from autoimmune diabetes

**DOI:** 10.1101/2023.07.19.549693

**Authors:** Shuaishuai Zhu, Emmanuelle Waeckel-Énée, Anna Moser, Marie-Andrée Bessard, Kevin Roger, Joanna Lipecka, Ayse Yilmaz, Barbara Bertocci, Julien Diana, Benjamin Saintpierre, Ida Chiara Guerrera, Stefania Francesconi, François-Xavier Mauvais, Peter van Endert

## Abstract

Appropriate tuning of protein homeostasis through mobilization of the unfolded protein response (UPR) is key to the capacity of pancreatic beta cells to cope with highly variable demand for insulin synthesis. An efficient UPR ensures a sufficient beta cell mass and secretory output but can also affect beta cell resilience to autoimmune aggression. The factors regulating protein homeostasis in the face of metabolic and immune challenges are insufficiently understood. We examined beta cell adaptation to stress in mice deficient for insulin-degrading enzyme (IDE), a ubiquitous protease with high affinity for insulin and genetic association with type 2 diabetes. IDE deficiency induced a low-level UPR in both C57BL/6 and autoimmune non-obese diabetic (NOD) mice, associated with rapamycin-sensitive beta cell proliferation strongly enhanced by proteotoxic stress. Moreover, in NOD mice, IDE deficiency protected from spontaneous diabetes and triggered an additional independent pathway, conditional on the presence of islet inflammation but inhibited by proteotoxic stress, highlighted by strong upregulation of regenerating islet-derived protein 2, a protein attenuating autoimmune inflammation. Our findings establish a key role of IDE in islet cell protein homeostasis, identify a link between low-level UPR and proliferation, and reveal an UPR-independent anti-inflammatory islet cell response uncovered in the absence of IDE of potential interest in autoimmune diabetes.

## 1. INTRODUCTION

Diabetes is the consequence of the failure of pancreatic beta cells to provide sufficient insulin to maintain normal blood glucose levels, due either to insulin resistance of tissue cells or to beta cell demise through an autoimmune response [1]. Physiologically, a sufficient insulin output is maintained by adaptive mechanisms including upregulation of beta cell secretory capacity but also beta cell proliferation and hyperplasia, together with protection from autoimmunity by regulatory T cells. Beta cell adaptation requires the mTORC1 (mammalian target of rapamycin 1) complex to activate and sustain protein translation, and the effectors of the UPR, a set of cellular pathways enabling the cell to cope with high protein loads in the endoplasmic reticulum (ER) by selective limitation of protein translation, increased production of chaperones, and upregulation of ER-associated protein degradation (ERAD) [2]. While both mTORC1 and the UPR are critical for maintaining a beta cell mass adapted to metabolic needs, chronic and/or excessive activation of each of these interconnected pathways can result in beta cell failure and ultimately death [3]. However, the factors affecting the setpoints between beneficial and deleterious UPR and mTORC1 activation are not completely understood. Moreover, while it is now recognized that an efficient UPR contributes to beta cell resilience in the face of autoimmune aggression [4], the interplay between beta cell physiology and autoimmunity remains insufficiently studied.

IDE is a ubiquitous endo-metalloprotease with high insulin affinity preferring small (≤ 10 kDa) substrates including some amyloidogenic peptides [5–7]. First identified as major insulin-degrading activity in hepatocytes, the role of IDE in physiologic insulin degradation remains unclear, given that mice with global or liver-restricted IDE deficiency display only mild hyperinsulinemia [8–10], and that endosomal insulin degradation by cathepsins has been described [11]. The wide expression in cells lacking insulin receptors as well as its high evolutionary conservation suggest that IDE may fulfill important functions beyond insulin degradation. Various lines of research have suggested a role of IDE in cellular protein homeostasis. IDE interacts both with proteasome complexes and ubiquitin [12,13], with a putative bimodal effect on proteasome activity [14]. IDE, which is induced by heat shock, has also been proposed to act as “dead-end chaperone” [15][6], for example forming complexes with aggregation-prone alpha-synuclein oligomers in beta cells [16] and with unfolded viral protein precursors [17], or degrading beta amyloid in a parallel ERAD pathway [18].

Moreover, knockout of a fission yeast IDE homologue increases resistance to tunicamycin, a drug inducing a strong UPR, in a TORC1-dependent manner, suggesting a role in cellular stress responses [19]. *Ide* polymorphism is associated with fasting glucose levels and the risk of human type 2 diabetes [20]. Global, liver- and beta cell-restricted IDE deficiencies in C57BL/6 mice induce pronounced to mild glucose intolerance [8–10,21]. Conflicting results were obtained with inhibitors of the IDE active or regulatory sites, ranging from ameliorated to acutely compromised glucose tolerance [22,23]. Thus, IDE expression both in beta and liver cells moderately affects glucose metabolism, but how its role can be manipulated to ameliorate it remains unclear.

Prompted by evidence for a role of IDE in protein homeostasis, here we studied C57BL/6 mice and NOD mice spontaneously developing autoimmune diabetes for activation of the UPR, inflammation and beta cell proliferation at the steady state and under metabolic, auto-immune or pharmacologic proteotoxic stress. We report that IDE deficiency protects NOD mice from autoimmune diabetes. Mechanistic studies reveal induction of a low-level UPR in both mouse strains associated with rapamycin-sensitive islet and beta cell proliferation. In addition, our findings suggest that auto-immune infiltration of *Ide^-/-^* NOD islets specifically induces strong upregulation of REG2 (regenerating islet-derived protein 2), a protein known to attenuate inflammation and promoting proliferation, and potentially responsible for reduced expression of some inflammatory cytokines and protection from diabetes.

## 2. RESULTS

### 2.1. IDE-deficient NOD mice display beta cell hyperplasia/hypertrophy and dysfunction combined with protection from autoimmune diabetes

To produce *Ide^-/-^* NOD mice, we back-crossed previously published *Ide^-/-^* C57BL/6 mice [8] to NOD mice (Figure S1). *Ide^-/-^* NOD mice had normal fasting glucose and insulin levels (Figure 1A, B). In oral glucose tolerance tests (OGTT), *Ide^-/-^* NOD mice displayed slightly increased glycemia at several time points, associated with hyperinsulinemia, consistent with the findings of Farris and our previous findings [23]. Normoglycemic female *Ide^-/-^* NOD mice had significantly increased numbers of islets relative to WT mice (Figure 1C) which secreted higher amounts of insulin upon glucose stimulation *in vitro* (Figure 1D). Moreover, islets from 8-week-old *Ide^-/-^* mice had a larger volume, although the difference to WT islets reached significance only for C57BL/6 and not for NOD mice (Figure 1E). Serum levels of islet amyloid polypeptide (IAPP), a well-known IDE substrate that is co-secreted with insulin and reduces post-prandial peaks of glycemia (24,25), were strongly increased in *Ide^-/-^* NOD mice aged 5 to 18 weeks (Figure 1F), presumably due to the absence of IAPP degradation by IDE. Collectively, these results are consistent with a role of IDE in regulating insulin and IAPP levels in beta cells and suggest that *Ide^-/-^* mice harbor an increased islet and beta cell mass which may result from beta cell hyperplasia, hypertrophy and/or regeneration, combined with compromised glucose tolerance and glucose-induced hyper-insulinemia.

**Figure 1:**
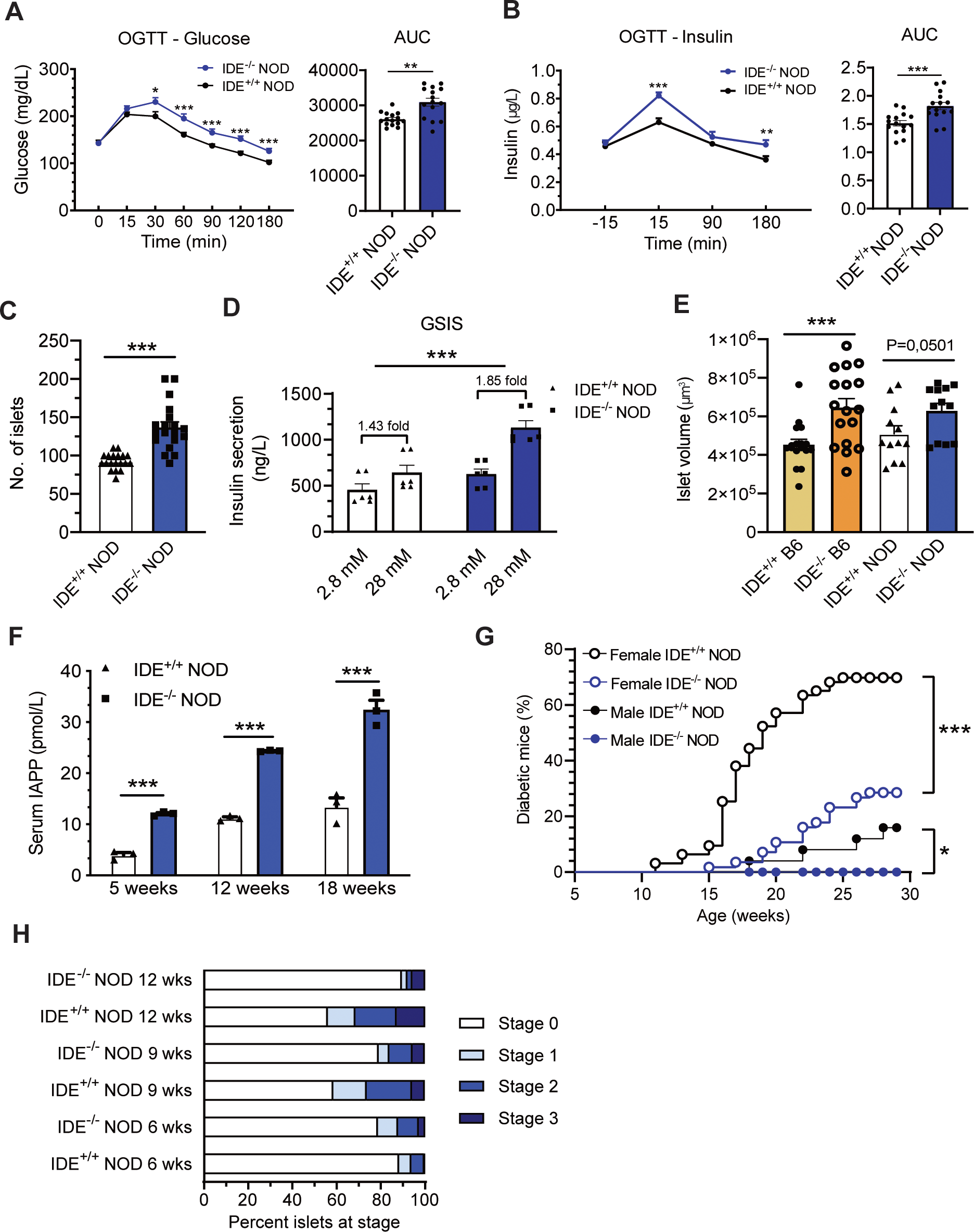
Beta cell function in *Ide^-/-^* C57BL/6J and NOD mice. **(A)** *Ide^+/+^* and *Ide^-/-^* NOD mice aged 14 weeks were subjected to an OGTT and glycemia was measured at indicated time points before (0 min) or after the glucose bolus. Data are mean values ± SEM of 15 independent mice for each group. **(B)** Insulinemia during the OGTT as measured by ELISA. Data are mean values ± SEM of 15 independent mice for each group. **(C)** Hand-picked islets obtained from female *Ide^+/+^* and *Ide^-/-^* NOD mice aged 8 weeks were counted in three separate experiments. Data are mean values ± SEM of 19 independent mice for each group. **(D)** Following 90 min incubation without glucose, hand-picked islets from 10-week-old *Ide^-/-^* and *Ide^+/+^* NOD mice were sequentially incubated for 1 h in 2.8- and 28-mM glucose, and insulin secreted into the supernatant was quantified by commercial ELISA. Data are mean values ± SEM of 6 mice for each group. **(E)** The volume of hand-picked islets from female C57BL/6 and NOD mice aged 8 weeks was calculated with Icy Software. Data are mean values ± SEM of an average of 15 mice for each group. Data in **(A-E)** were evaluated by Student’s t-test, *p < 0.05, **p < 0.01, ***p < 0.001. **(F)** The amount of IAPP in the serum of female mice of different age was quantified by ELISA. Data are mean values ± SEM of 4 independent mice for each group. ***p < 0.001 by Student’s t-test. **(G)** *Ide^+/+^* NOD and back-cross 10 *Ide^-/-^* NOD mice were monitored weekly for glycosuria until 29 weeks of age. Data are mean values ± SEM. N = 63 for female *Ide^+/+^*, 57 for female *Ide^-/-^*, 24 for male *Ide^+/+^*, 29 for male *Ide^-/-^* mice. ***p < 0.001 by Kaplan-Meier, log-rank (Mantel-Cox) test. **(H)** Islets of pre-diabetic female *Ide^+/+^* and *Ide^-/-^* mice of different age were stained with hematoxylin/eosin and scored for insulitis. Score 0, no infiltration; 1, peri-insulitis; 2, moderate intra-insulitis (<50% of islet surface); 3, severe insulitis (>50 of surface and/or loss of islet architecture). Between 190 and 350 islets were counted for each group.

Wondering how this beta cell dysfunction might affect manifestation of autoimmune diabetes, we monitored glycemia in *Ide^-/-^* NOD mice. IDE deficiency provided significant though incomplete protection from T1D to mice of both sexes, with 28% of *Ide^-/-^* vs. 70% of *Ide*^+/+^ female (p<0.0001), and 0% of *Ide^-/-^* vs. 17% of *Ide*^+/+^ male (p<0.05) animals manifesting hyperglycemia by 30 weeks (Figure 1G). This was associated with a decrease in islet inflammation between 6 and 12 weeks of age, contrasting with the well-known increased insulitis (stage 2 or 3) in *Ide*^+/+^ NOD mice (Figure 1H).

### 2.2. IDE deficiency induces distinct stress responses in NOD and C57BL/6 pancreatic islets

Although T2D eventually results in a reduced beta cell mass, an increased beta cell and insulin mass can develop to compensate for high metabolic demand in obesity, insulin resistance or due to beta cell-intrinsic factors [24–26]. A main driver for compensatory upregulation of beta cell mass and function is the UPR, which is constitutively upregulated in beta cells [27][28]. Given our finding of an increased islet cell mass in *Ide^-/-^* islets (Figure 1) and literature suggestions of a role in protein homeostasis as a chaperone and regulator of the proteasome-ubiquitin system, we asked whether *Ide^-/-^* islets displayed evidence of UPR upregulation.

We first tested fresh handpicked islets from pre-diabetic NOD mice and control C57BL/6 mice for expression of genes representing the three branches of the UPR, spliced X-Box binding protein 1 (XBP1s) (indicating inositol-requiring enzyme 1alpha (IRE1α) activation), activating transcription factor 6 (ATF6) and C/EBP homologous protein (CHOP, a pro-apoptotic protein downstream of protein kinase RNA-like endoplasmic reticulum kinase (PERK)). We also included immunoglobulin-binding protein (BIP, GRP78), an ER chaperone with a key role in the UPR, proliferating cell nuclear antigen (PCNA) and Ki67 as markers of proliferation, and regenerating islet-derived 2 (REG2), a protein expressed in regenerating islets and but not in normal islets [29,30]. REG2 has been reported to increase the beta cell mass, ameliorate diabetes in NOD mice and attenuate the UPR and apoptosis [31–33].

Islets from female *Ide^-/-^* C57BL/6 mice aged 7 weeks displayed strong upregulation of BIP mRNA but not of other UPR or proliferation markers. This upregulation was maintained after overnight culture of islets, suggesting that BIP induction was due to an islet-intrinsic mechanism (Figure 2A-C). In striking contrast, islets from *Ide^-/-^* female NOD mice aged 7 or 12 weeks lacked BIP induction but displayed strong REG2 upregulation (Figure 2D, E). However, this was completely lost upon overnight culture (Figure 2F), a procedure resulting in rapid exit of infiltrating immune cells from islets [34]. Therefore, upregulation of REG2 may be triggered by autoimmune infiltration. Supporting this conclusion, islets devoid of inflammation from *Ide^-/-^ Rag2^-/^*^-^ mice completely lacked REG2 induction, and islets from male *Ide^-/-^* mice with attenuated infiltration displayed lower REG2 induction that was also rapidly lost *ex vivo* (Figure 2G, H, S2A, B). This was not due to absence of REG2 degradation by IDE which was unable to digest REG2 even upon prolonged *in vitro* incubations (Figure S2C, D). Taken together, these results suggested that the absence of IDE induces a mild UPR in C57BL/6 islets, consistent with its role in protein homeostasis. Conversely *Ide^-/-^* NOD islet cells upregulate a key marker of regeneration in the presence of inflammation but fail to upregulate BIP, possibly in relation to a defective UPR reported for this strain [4].

**Figure 2:**
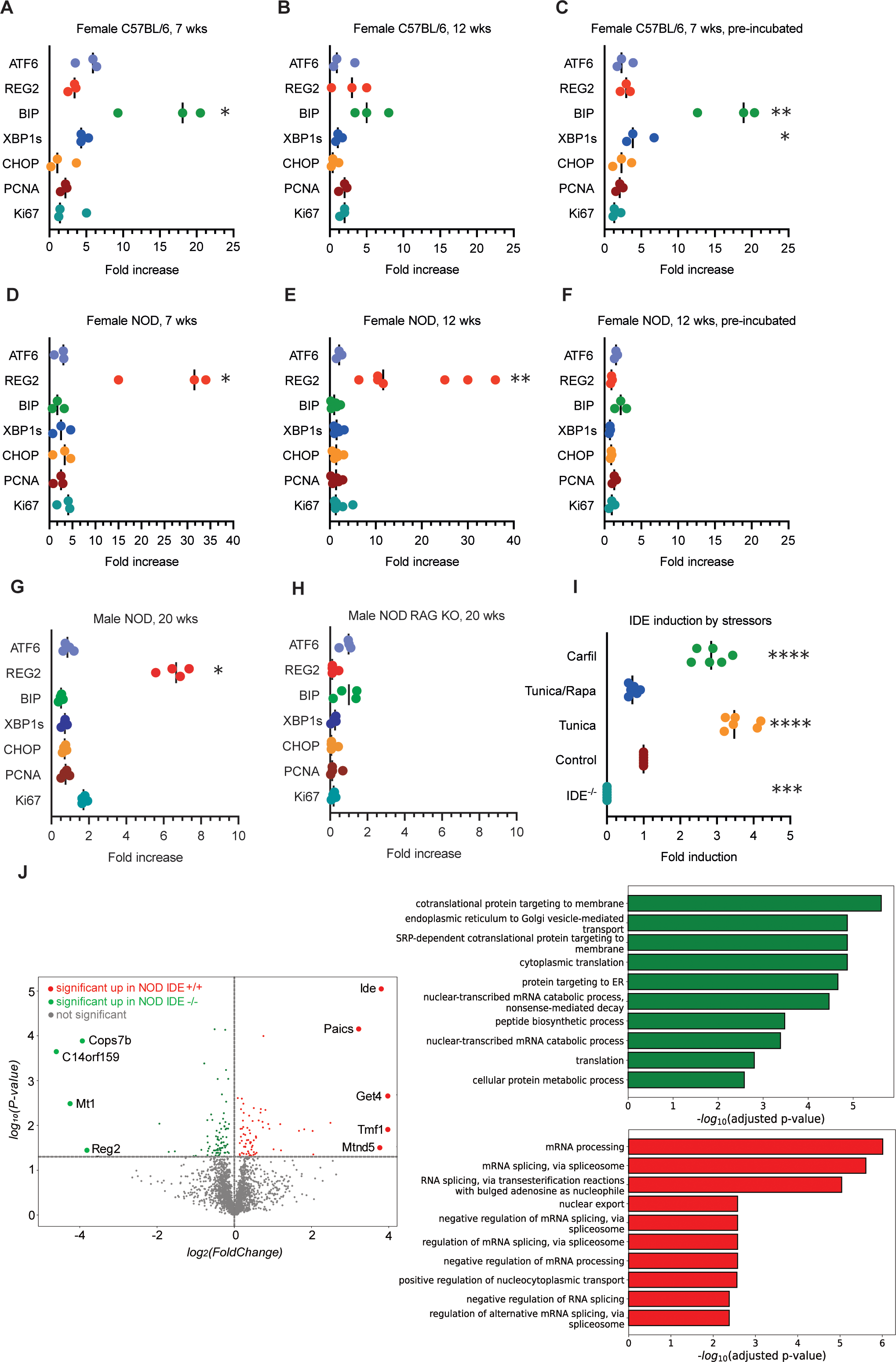
UPR effector mRNA and global protein expression by IDE^-/-^ islet cells. **(A-H)**, mRNA expression levels, as measured by RT-qPCR, of genes linked to the UPR (ATF6, BIP, XBP1s, CHOP), to regeneration (REG2) or to proliferation (PCNA, Ki67) in steady state islets from *Ide^+/+^* and *Ide^-/-^* C57BL/6 and NOD mice of different ages, as indicated. Results are expressed as fold change, i.e., the ratio of expression in *Ide^-/-^* relative to *Ide^+/+^* islets. Islets were either analyzed immediately after isolation or after overnight culture in RPMI medium, as indicated. Each dot represents islets from one mouse. Data in **(A-H)** were evaluated by Student’s t-test. p < 0.05, **p < 0.01, ***p < 0.001, ****p < 0.0001. **(I)** *Ide^+/+^* or control *Ide^-/-^* NOD mice aged 10 weeks were injected for 2 weeks with carfilzomib or for 2 days with tunicamycin with or without addition of rapamycin before isolation of islets and quantification of IDE mRNA expression by RT-qPCR. **(J**) Proteomic analysis of islet proteins from 3 *Ide^+/+^* and 3 *Ide^-/-^* mice aged 10 weeks. Left: volcano plot showing the proteins up-(in green) and down-(in red) regulated in *Ide^-/-^* NOD islets. The horizontal dashed line represents the significance threshold (p-value < 0.05) (for details see supplementary table 1). Right: Barplot representing the Gene Ontology Biological Process (GO BP) terms enriched using enrichR software and the GO Biological Process 2021 database. In green are the TOP10 GO BP terms enriched using the up-regulated proteins in *Ide^-/-^* and in red the TOP10 GO BP terms enriched using the up-regulated proteins in *Ide^+/+^* islets for analysis. A log-transformed adjusted p-value was used for TOP10 ranking (for details see supplementary table 2).

To obtain additional evidence for a role of IDE in islet cell protein homeostasis, we treated NOD mice for two days with a proteasome inhibitor or tunicamycin and examined IDE mRNA levels by islets. IDE expression was strongly upregulated by both drugs, consistent with a role in the response to proteotoxic stress (Figure 2I). Interestingly, mTORC1 inhibition by rapamycin abolished the effect of tunicamycin on IDE expression, suggesting that IDE levels may correlate with the global level of protein translation in stressed cells.

To obtain a broad view on the effect of IDE deficiency on islet protein expression, we subjected lysates of *Ide^+/+^* and *Ide^-/-^* NOD islets to a global proteomic analysis. Analysis of protein abundance revealed that both WT and *Ide^-/-^* cells were enriched for pathways related to mRNA processing, however these concerned nuclear RNA export and splicing in WT versus RNA degradation in *Ide^-/-^* cells (Figure 2J). Proteins in *Ide^-/-^* islets were also enriched for translation of ER-targeted proteins and ER to Golgi protein transport (Figure 2J and Tables S1, S2), consistent with upregulation of the UPR. Differentially expressed genes in WT cells included the BAG6/BAT3 complex (Get4), a platform for sorting of proteins between the ER and cytosolic degradation, and Tmf1 (TATA element modulatory factor), a protein essential for Glut4 trafficking to Glut4 storage vesicles. *Tmf^-/-^* mice develop hyperglycemia due to compromised glucose uptake [35]. Proteins upregulated in *Ide^-/-^* cells included the proteasome-associated COP9 signalosome (Cops7b), which decreases protein ubiquitylation by SCF-type E3 ligases, metallothionin (Mt1), involved in zinc ion homeostasis and upregulated in beta cell stress, and importantly REG2 (Figure 2J). Collectively these results are consistent with a role of IDE of protein homeostasis by affecting regulation of transcription, translation, secretory pathway transport and degradation of proteins.

To obtain additional insight on the relationship between IDE expression and protein homeostasis, we next analyzed islets obtained from mice treated with tunicamycin which induces strong ER stress. *Ide^-/-^* C57BL/6 islets responded with stronger upregulation of BIP to this treatment than WT islets while other stress or proliferation marker mRNAs were little or not affected in both WT and *Ide^-/-^* islets (Figure S3A-C). In WT NOD islets, tunicamycin induced a moderate upregulation of all markers including REG2 except for ATF6 (Figure S3D-F). In contrast, *Ide^-/-^* islets displayed induction of BIP, XBP1s and Ki67 whereas REG2 was down-regulated (Figure S3G-I). Thus, REG2 is upregulated by islet inflammation but downregulated by massive ER stress in *Ide^-/-^*NOD islets.

We also examined ER stress signaling at the protein level. At the steady state, PERK and eIF2α phosphorylation, ATF6 nuclear translocation and PCNA expression did not differ between WT and *Ide^-/-^* C57BL/6 mice (Figure 3A-G). Tunicamycin significantly induced all three ER stress effectors and PCNA in *Ide^-/-^* islets, an effect abolished by mTORC1 inhibition, while only ATF6 translocation was triggered in WT islets (Figure 3B-G). Thus, C57BL/6 *Ide^-/-^* islet cells display increased sensitivity to massive proteotoxic stress. In contrast to C57BL/6 islets, steady state islets from *Ide^-/-^* NOD mice aged 9-15 weeks were characterized by a partial stress response absent in WT islets, with phosphorylated eIF2α but not PERK, increased nuclear ATF6 and upregulated PCNA (Figure 3H-N). Thus, islets from *Ide^-/-^* NOD mice displayed a specific partial UPR next to REG2 mRNA induction which may be due to their genetic background and/or to the presence of autoimmune inflammation, the latter being consistent with the results shown in Figure 2G, H.

**Figure 3.**
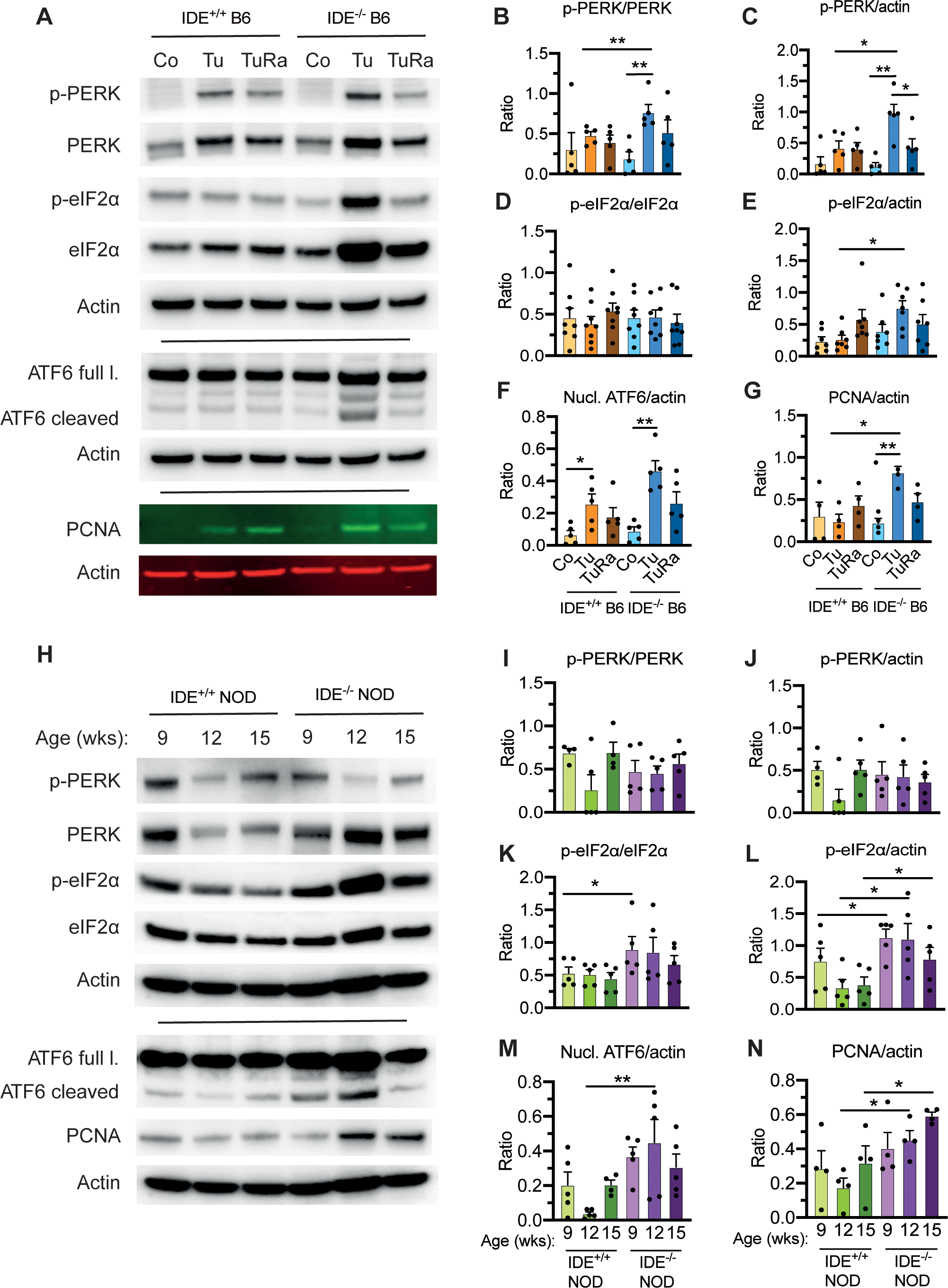
Immunoblot analysis of UPR effector activation. **(A)** Representative immunoblots showing the expression pattern of phosphorylated PERK, total PERK, phosphorylated eIF2α, total eIF2α, ATF6, cleaved ATF6, and PCNA in islet lysates of C57BL/6 *Ide^+/+^* and *Ide^-/-^* mice treated by tunicamycin injection i.p., combined with rapamycin or not, for 48 hours. **(B-G)** quantification of experiments performed as shown in **(A)**. N = 5 in **(B)**, 5 in **(C)**, 8 in **(D)**, 7 in **(E)**, 5 in **(F)** and 4 in **(G)**. Data represent the ratio of *Ide^-/-^* to *Ide^+/+^* islets, are represented as mean ± SEM and were evaluated by Student’s t-test in **(B-F)** and by ANOVA in **G**. *P < 0.05, **P < 0.02. **(H)** Representative protein immunoblots as in **(A)** but for steady state islets from *Ide^+/+^* and *Ide^-/-^* NOD mice aged 9, 12 or 15 weeks. **(I-N)** quantification of experiments performed as shown in **(H)**. N = 5 for panels **(I-M)** and 4 for **(N)**. Data are represented as mean ± SEM and were evaluated by Student’s t test; *p < 0.05.

### 2.3. IDE deficiency modulates inflammatory signaling differentially in C57BL/6 versus NOD mice

Considering that ER stress and insulin signaling can induce an inflammatory response [36], we next examined islets for expression of inflammatory marker mRNA. Steady state islets from young (aged 4 to 12 weeks) female *Ide^-/-^* NOD mice displayed downregulation of IL-1β mRNA and strong down-regulation of 2’-5’-oligoadenylate synthase 3 (OAS-3), a key enzyme in type 1 interferon signaling while expression of additional inflammatory genes was not altered (Figure 4, S4A). MRNA expression of a subset of markers in male NOD islets was not altered (Figure S4B). After short-term tunicamycin treatment of female NOD mice, OAS-3 expression remained low while expression of some inflammatory markers (NLRP3, MCP-1, IF27L2A) was moderately increased (Figure 4B).

**Figure 4:**
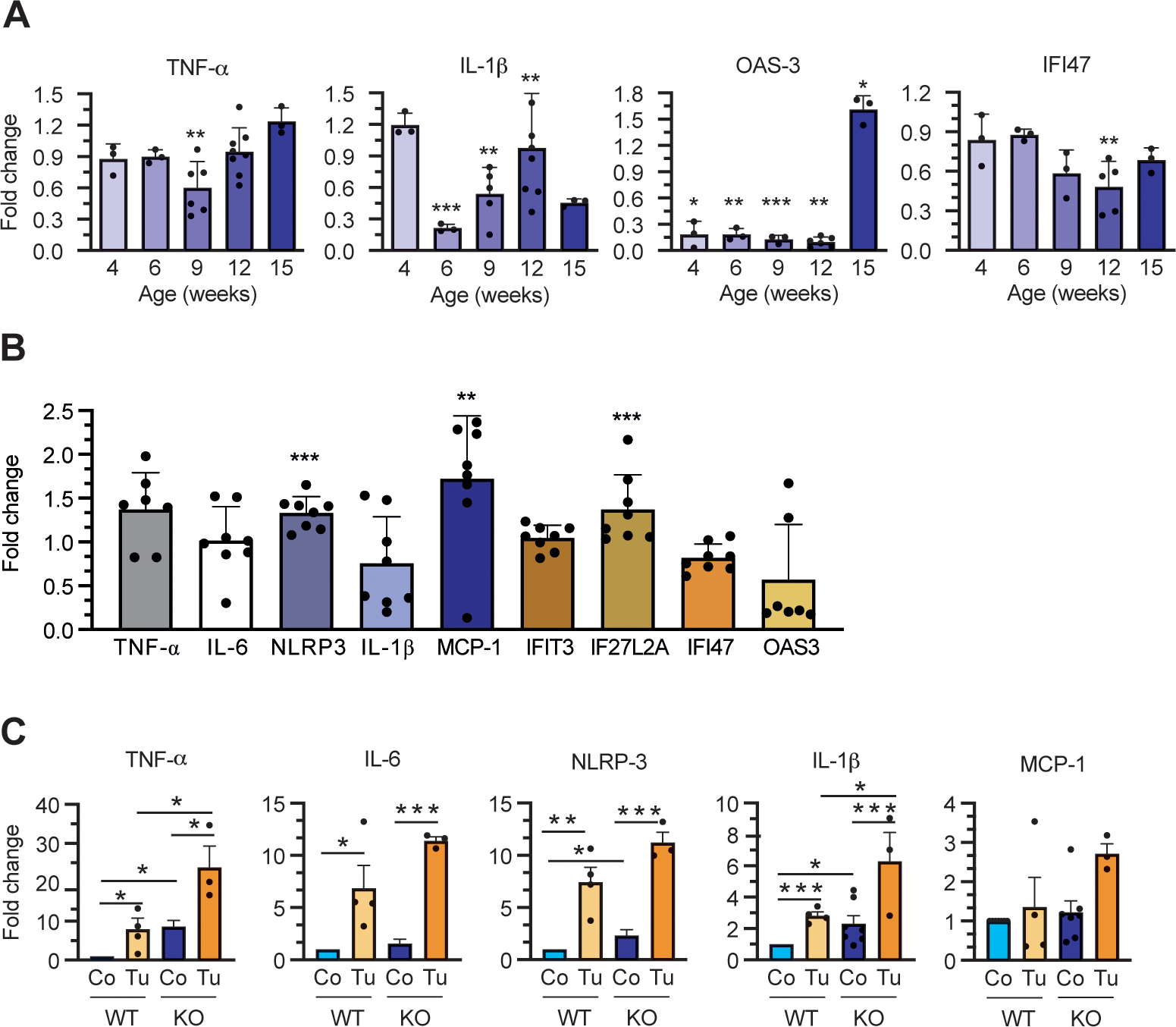
Effect of IDE deficiency on expression of inflammatory genes. **(A)** RT-qPCR analysis of relative TNF-α, IL-1β, OAS-3, and IFI47 expression in islets of female *Ide^+/+^* and *Ide^-/-^* NOD mice aged 4 to 15 weeks. Results represent the ratio of expression in *Ide* ^-/-^ relative to *Ide* ^+/+^ islets and are mean values ± SEM of 3-6 independent mice for each group. *P < 0.05, **P < 0.02, and ***P < 0.01 by Student’s t-test. **(B)** mRNA expression of the indicated genes in the islets of *Ide^-/-^* relative to *Ide^+/+^* NOD mice treated with tunicamycin for 48 hours. Data are mean values ± SEM of 8 independent mice for each group. *p < 0.05, **p < 0.02, and ***p < 0.01 by Student’s t-test for panels **(A, B)**. **(C)** mRNA expression of the indicated genes in *Ide^-/-^* relative to *Ide^+/+^* islets from C57BL/6 mice aged 15 to 25 weeks treated with tunicamycin for 48 hours. Data are mean values ± SEM of 6 independent mice for each group. *p < 0.05, **p < 0.02, and ***p < 0.01 by ANOVA test.

Contrasting with NOD islets, expression of TNF-α, NLRP3 and IL-1β mRNA was increased in steady state islets from *Ide^-/-^* C57BL/6 as compared to WT mice. Tunicamycin treatment strongly increased expression of these and IL-6 mRNA in WT and *Ide^-/-^* islets, with expression by *Ide^-/-^* islets remaining superior (Figure 4C). Thus, IDE deficiency had a markedly different effect on transcription of inflammatory effectors, with significant to strong activation in *Ide^-/-^* C57BL/6 islets contrasting with downregulation in *Ide^-/-^* NOD islets.

### 2.4. High fat diet induces obesity and decompensation of glucose metabolism in *Ide^-/-^*mice

Considering our observations of compromised glucose tolerance and islet hyperplasia in *Ide^-/-^* mice (Figure 1), we subjected WT and *Ide^-/-^* C57BL/6 mice to high fat diet (HFD) for 7 weeks and monitored glucose metabolism and ER stress markers. In *Ide^-/-^* mice, HFD resulted in a massive weight gain by almost 70%, contrasting with 30% in WT mice; *Ide^-/-^* mice on standard diet were not overweight (Figure 5A, B). This was accompanied by massive hyperinsulinemia and even stronger hyperproinsulinemia in *Ide^-/-^* mice fed HFD (Figure 5C, D). HFD increased the amount of insulin and proinsulin in WT islets about 6-fold and in *Ide^-/-^*mice 10-fold (Figure 5 E, F); however, despite this massive upregulation of (pro-)insulin production, WT mice developed moderate and *Ide^-/-^* mice substantial hyperglycemia (Figure 5G).

**Figure 5:**
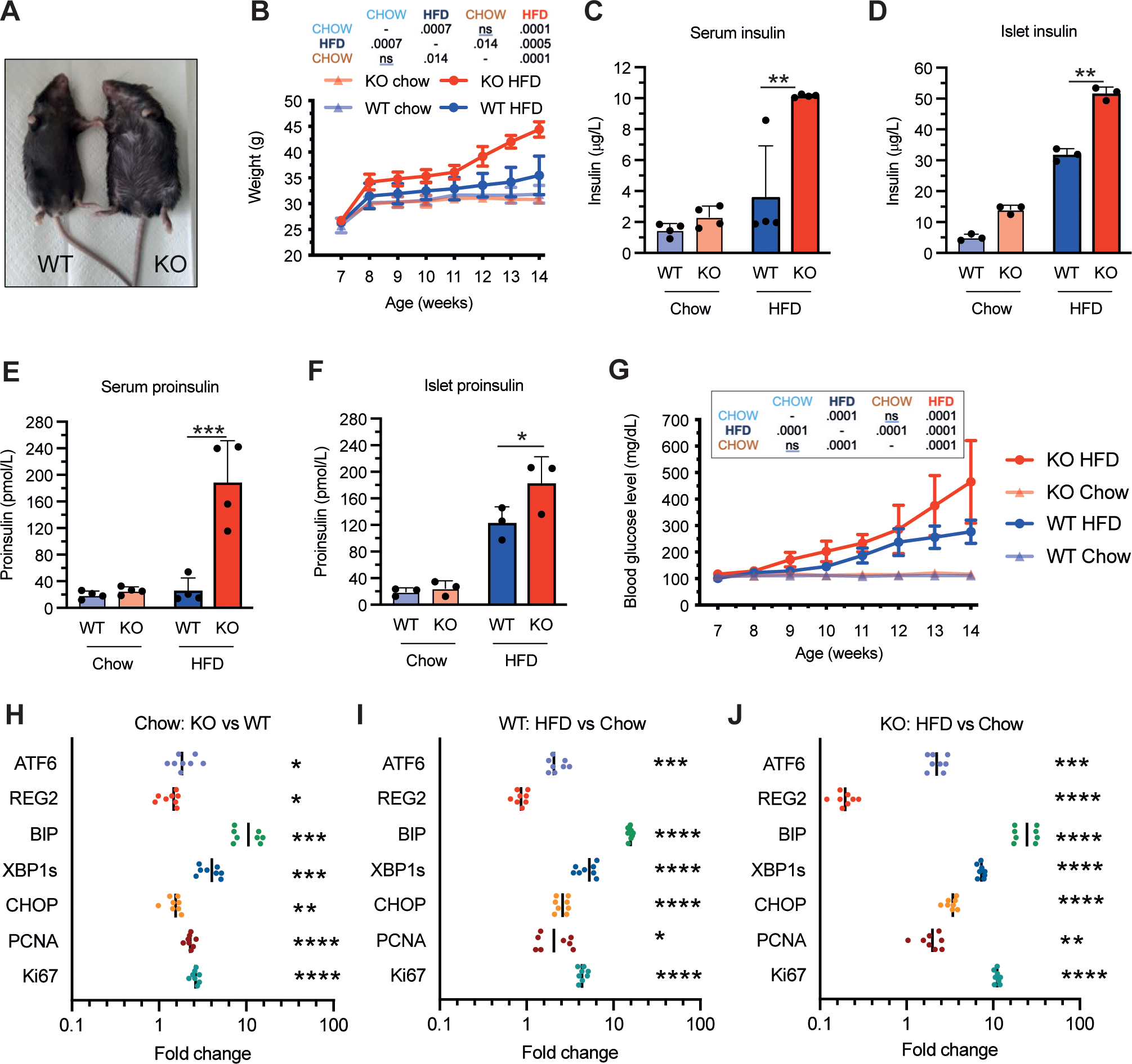
Effect of IDE deficiency on the metabolic and UPR response to HFD. Seven-week-old female *Ide^-/-^* and *Ide^+/+^* C57BL/6J mice were fed a standard chow or HFD for 7 weeks. N = 10 per group. **(A)** Representative appearance of *Ide^+/+^* and *Ide^-/-^* mice at sacrifice. **(B)** Body weight was recorded once a week. Data are mean values ± SEM. Statistical significance is indicated in the insert. **(C-F)** Serum insulin and proinsulin levels and total insulin and proinsulin content in islets were determined for *Ide^-/-^* and *Ide^+/+^* mice at the time of sacrifice, N = 6 per group. **(G)** Non-fasting glucose concentrations were monitored weekly; N = 10 per group. **(H-J)** Expression levels of genes related to ER stress, proliferation, and regeneration, as measured by RT-qPCR, in islets obtained at sacrifice of *Ide^-/-^* and *Ide^+/+^* mice fed standard chow or HFD, N = 8. Results are expressed as ratio of expression as indicated: *Ide^-/-^* vs. *Ide^+/+^* in standard chow-fed **(H)**, HFD vs. standard chow for *Ide^+/+^* mice **(I)** or for *Ide^-/-^* mice **(J)**. All panels show mean values ± SEM, plus individual data points in **(H-J)**. Statistical evaluation was performed by two-way ANOVA with Sidac correction for multiple comparisons in panels **(B, G)** and by Student’s t-test for all other panels. *p < 0.05, **p < 0.01, ***p < 0.001, ****p < 0.0001.

Next, we examined the effect of metabolic stress on UPR and proliferation markers. Islets from *Ide^-/-^* C57BL/6 mice fed standard diet showed prominent induction of BIP like at steady state. HFD resulted in upregulation of all markers both in WT and *Ide^-/-^* islets except for REG2 the expression of which was not affected in WT but down-regulated in *Ide^-/-^* islets (Figure 5H-J). Next to BIP, Ki67 expression was most strongly affected by HFD and induced 10-fold in *Ide^-/-^* islets (Figure 5I, J). Thus, *Ide^-/-^* islets react to extreme metabolic demand by amplifying the response already present in WT islets; interestingly, this response includes Ki67 suggestive of cell proliferation but not REG2 which is mobilized specifically by immune stress in *Ide^-/-^* NOD islets.

Given the association of increased (pro-)insulin output by beta cells with UPR induction, we wondered whether conversely inducing a UPR can induce insulin output by islet cells. Indeed, tunicamycin treatment induced *in vivo* secretion of insulin and, to a lesser extent, proinsulin in both WT and *Ide^-/-^*NOD mice, which was abolished by the mTORC1 inhibitor rapamycin (Figure S5A, B). However, while *Ide^-/-^* islets contained higher amounts of insulin and proinsulin than WT islets, short-term *in vivo* treatment by the drugs had no significant effect on these amounts (Figure S5C, D).

### 2.5. IDE deficiency induces beta cell proliferation enhanced by metabolic and pharmacologic stress

The UPR induces cellular adaptation through ER expansion but also through beta cell proliferation. We analyzed islets at the steady state and from mice exposed to metabolic or pharmacologic stress for expression of Ki67 and incorporation of 5-ethynyl-2′-deoxyuridine (EdU). At the steady state, NOD *Ide^-/-^* islets contained up to six-fold more Ki67 spots than WT islets. Tunicamycin upregulated Ki67 expression strongly in both WT and *Ide^-/-^* NOD islets, an effect abolished by rapamycin and by salubrinal, an inhibitor of eIF2α dephosphorylation (Figure 6A, B, C). Steady state *Ide^-/-^* C57BL/6 islets also displayed higher Ki67 staining than WT islets, a difference amplified in mice fed a HFD (Figure 6D, E). To directly measure DNA replication in beta cells, we compared EdU incorporation in *Ide^-/-^* insulin-positive cells. EdU incorporation was increased at the steady state both in C57BL/6 and NOD beta cells (Figure 6F-I). Thus, *Ide^-/-^* islet and beta cells display constitutive upregulation of proliferation; metabolic stress enhances proliferation in WT and *Ide^-/-^* cells in a manner dependent on mTORC1 and on eIF2α dephosphorylation but not REG2 induction.

**Figure 6:**
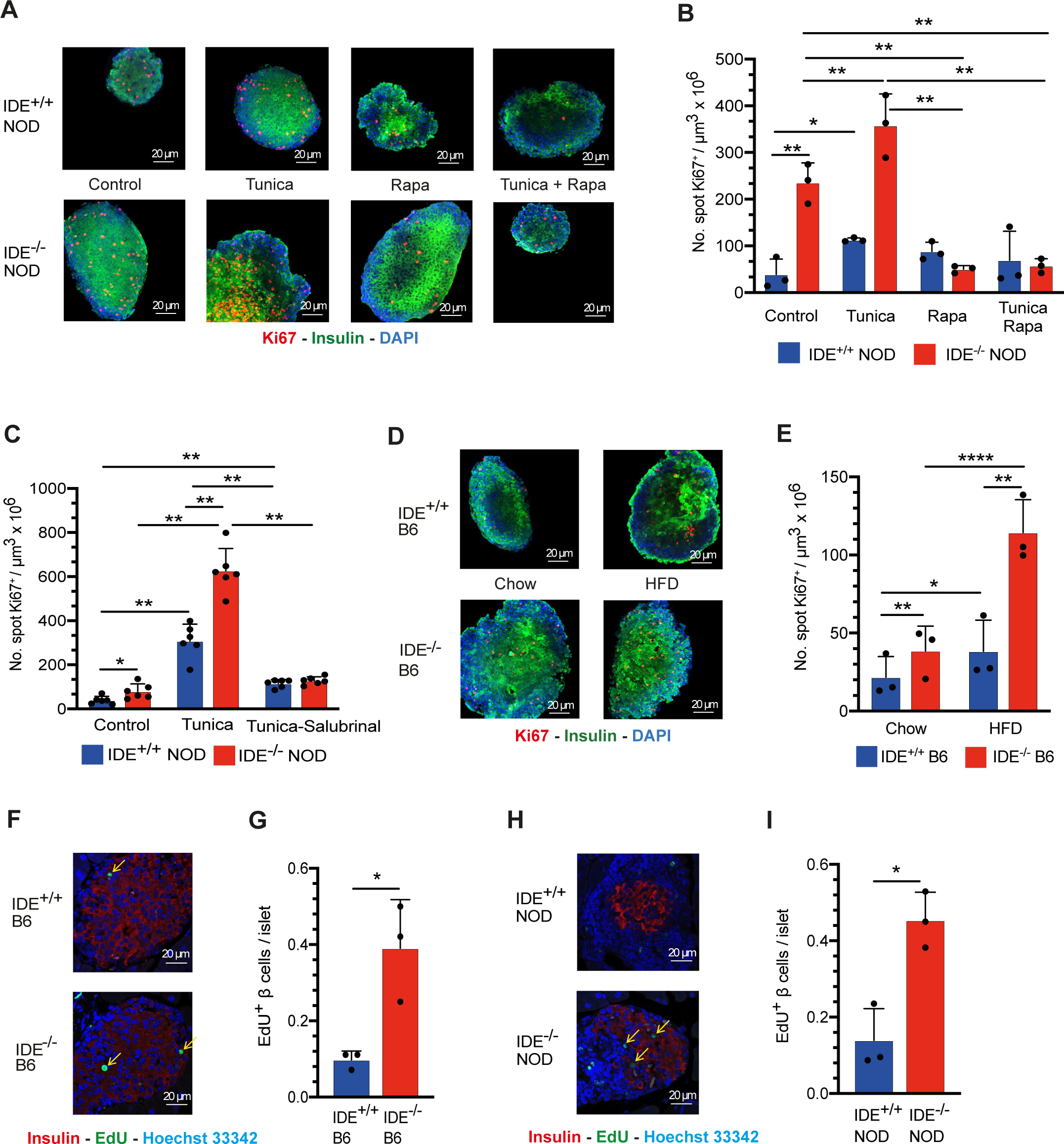
Effect of proteotoxic or metabolic stress and of IDE deficiency on islet cell proliferation. **(A)** Representative images stained for insulin (green) and Ki67 (red) of islets of female *Ide^-/-^* or *Ide^+/+^* NOD mice aged 12 weeks and treated for 2 days with tunicamycin and/or rapamycin or solvent alone. **(B)** Number of Ki67 spots per volume for the conditions indicated in **(A)**. N = 3 per group, with an average of 40 islets per mouse counted. **(C)** *Ide^-/-^* and *Ide^+/+^* NOD mice aged 12 weeks were treated for 2 days as in **(A)**, however using tunicamycin alone or combined with salubrinal. Quantitative evaluation as in **(B)**. N = 6 per group. **(D)** Representative images, stained for insulin and Ki67, showing islets from *Ide^-/-^* and *Ide^+/+^* C57BL/6 mice aged 14 weeks, after a HFD or standard diet for 7 weeks. **(E)** Quantitative evaluation as in **(B)** of the experiment shown in **(D)**. N = 3 per group. **(F)** Representative immunofluorescence images showing insulin expression and EdU incorporation in *Ide^-/-^* and *Ide^+/+^* C57BL/6 mice after treatment with tunicamycin for 2 days followed by administration of EdU for 1 h. The arrows point to EdU^+^ beta cells. **(G)** Quantification of the number of EdU^+^ beta cells per islet from the experiment shown in **(F)**. N = 3 mice per group, with on average 55 islets per mouse evaluated. **(H)** Representative immunofluorescence images showing insulin expression and EdU incorporation in steady state islets from *Ide^-/-^* and *Ide^+/+^* NOD mice aged 12 weeks. The arrows point to EdU^+^ beta cells. **(I)** Quantification of EdU^+^ beta cells in **(H)**. N = 3 mice per group, with on average 66 islets per mouse counted. The numbers of Ki67^+^ spots in panels **(B, C, E)** and the numbers of EdU^+^ beta cells in **(G, I)** are represented as mean ± SEM. Statistical analysis for all experiments was performed by Student’s t-test, *p < 0.05, **p < 0.01, ***p < 0.001, ****p < 0.0001.

2.6. RNAseq analysis confirms upregulation of *Ide^-/-^* islet cell proliferation upon HFD

To obtain initial mechanistic insight in the response of islet cells to ER stress, we performed RNAseq analysis on C57BL/6 mice subjected to HFD for 2 weeks. Gene expression among islets from 3 *Ide^-/-^* and 5 WT mice, respectively, correlated and *Ide^-/-^* islets clustered in principal component analysis while expression by WT mice was more heterogeneous (Figure 7A, B). Top genes enriched in WT islets were mainly related to protein metabolism and degradation but also included genes related to beta cell function: *Pnliprp1*, (inactive pancreatic lipase-related protein 1) and *Slc39a5* (zinc transporter ZIP5). In contrast, *Ide^-/-^* islets were strongly enriched for genes related to cycling, including *lockd* (long non-coding RNA regulating Cdkn1b), *Ckap2* (cytoskeleton-associated protein 2); and for genes related to stress responses: *Pyroxd2* (Pyridine-nucleotide disulfide oxidoreductase) and *Gsto2* (glutathione S-transferase omega-2) (Figure 7C).

**Figure 7:**
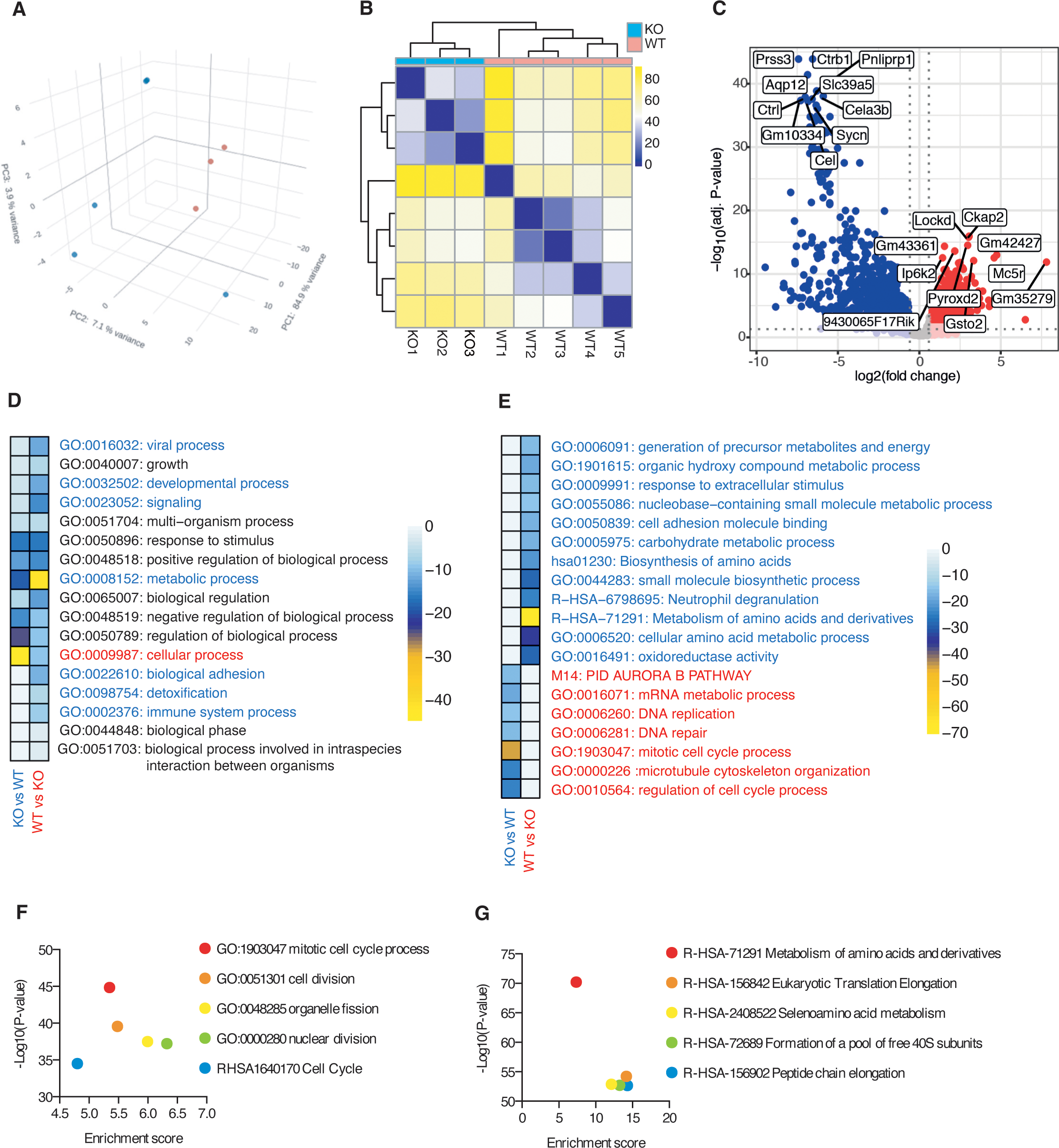
Gene expression in *Ide^-/-^* and *Ide^+/+^* islets of mice subjected to metabolic stress. **(A)** Relative distance between the bulk RNAseq transcriptome of individual *Ide^+/+^* and *Ide^-/-^* pancreatic islets purified from C57BL/6 mice exposed to HFD treatment for 14 days was assessed using principal component analysis. **(B)** Correlation matrix of *Ide^+/+^* and *Ide^-/-^* pancreatic islets based on all gene probes. **(C)** MA-plot of the log_2_ fold change (l2fc) of all genes. Blue and red dots indicate genes with P-value <0.05 enriched in *Ide^+/+^* and *Ide^-/-^*samples, respectively. The names of the top 10 discriminatory genes are indicated. **(D-G)** The lists of differentially expressed genes (adjusted p-value<0.01, l2fc>2) in *Ide^-/-^* and *Ide^+/+^* samples were compared using Metascape. Color heatmaps show the top parent GO biological processes **(D)** and the selected top GO biological process **(E)** Enriched pathways were calculated and clustered by adjusted p-values by automatic Metascape algorithm. Dot plots show the adjusted –Log_10_(p-value) and Enrichment score for the top 5 pathways specifically enriched in *Ide^-/-^* **(F)** and *Ide^+/+^* **(G)** samples. See also Figure S4.

Gene set enrichment analysis of differentially expressed genes indicated enrichment of cellular processes related to cell cycle, mitosis, cytokinesis and mRNA processing in *Ide^-/-^* islets from HFD-fed mice (Figure 7D, E; S6A); all five top gene sets were related to mitosis and cell cycle (Figure 7F). RNAseq analysis also confirmed that REG2 and additional tissue-protective antimicrobial peptides (REG3a, b, d) were not enriched in *Ide^-/-^* HFD islets. In contrast, genes differentially expressed in WT islets were strongly enriched in metabolic pathways related to amino acid metabolism and peptide chain elongation (Figure 7 D, E, G). Considering the link between UPR and inflammation and the enrichment in GO terms related to the immune system process (Figure 7D), we also analyzed immune-related gene sets. WT islets were enriched in genes related to antigen uptake (lectins, FcγR, C1q), antigen presentation (H-2Eβ) and processing (CD74, cathepsin S) (Figure S6B). Collectively, these data confirmed strong upregulation of proliferation in metabolically stressed C57BL/6 *Ide^-/-^* islets and suggested that IDE deficiency may reduce immune activation in this condition.

## 3. Discussion

In the context of islet cell biology, IDE has previously been studied essentially with respect to its role in glucose metabolism and insulin degradation. While we confirm previous findings such as compromised glucose tolerance and increased insulin content in *Ide^-/-^* islets, we have uncovered a role of IDE in islet protein homeostasis revealed by a basal and stress-enhanced activation of the UPR and consecutive mTORC1-dependent islet and beta cell proliferation. We also reveal a clearly distinct response to IDE deficiency in immune cell-infiltrated NOD islets associated with an attenuated inflammatory response, possibly related to the induction of REG2 expression. Thus, both at the steady state and upon massive proteotoxic stress, IDE deficiency provides the unexpected benefits of inducing C57BL/6 and NOD proliferation dependent on eIF2α dephosphorylation and mTORC1 but not REG2, whereas production of the anti-inflammatory molecule REG2 is induced only in inflamed NOD islets and abolished by massive proteotoxic stress (Figure 8).

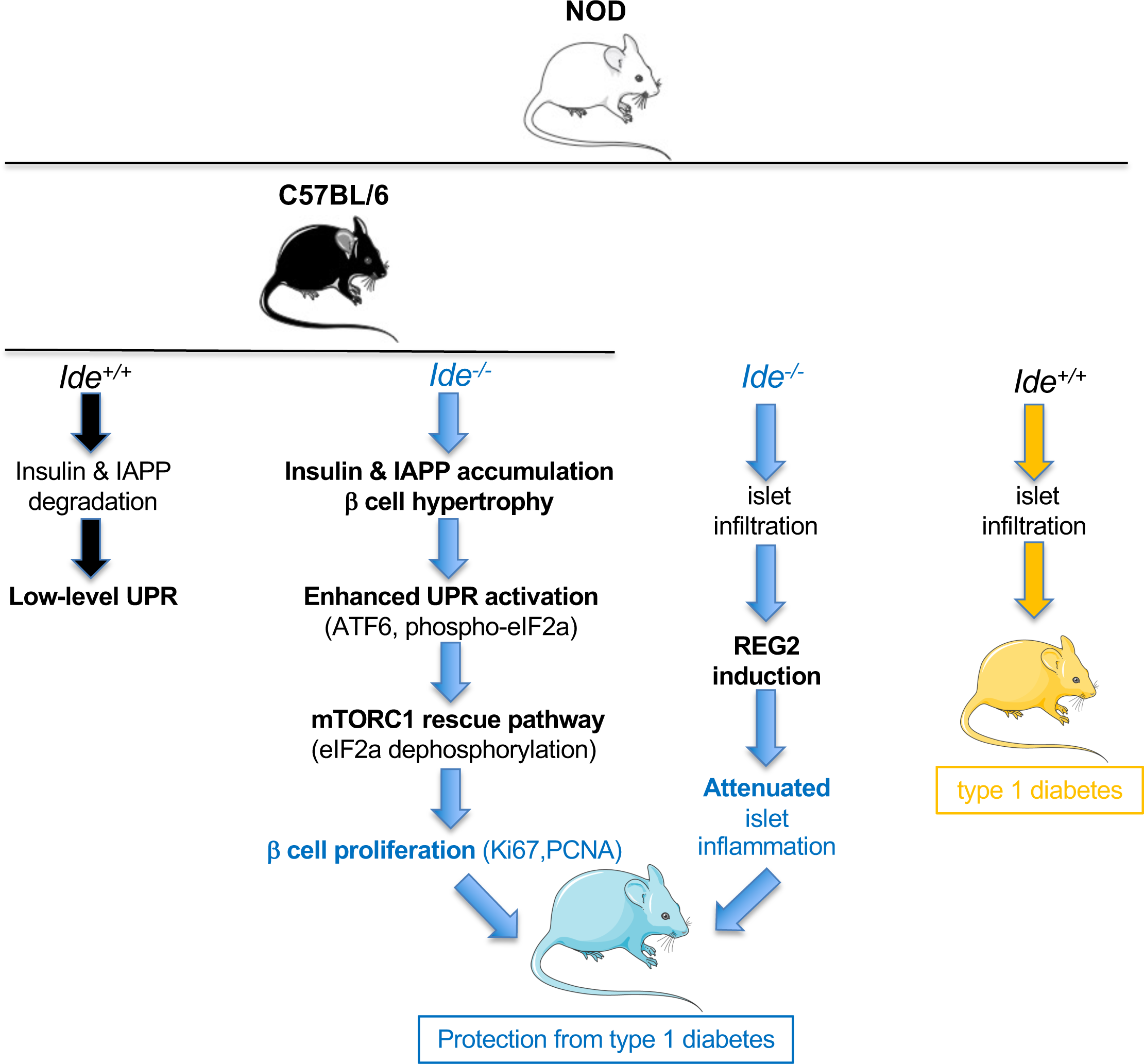

Although not previously described, a role of IDE in islet protein homeostasis is consistent, next to its high affinity for insulin, with evidence for interaction with, and regulation of, the ubiquitin-proteasome system, and its direct induction by proteotoxic stress (Figure 2I). Increased insulin and IAPP content of *Ide^-/-^* islets suggests a key role of IDE in degrading these substrates in beta cells. However, as folding of neither of these substrates is affected by inhibition of N-glycosylation, UPR enhancement by tunicamycin indicates that other substrates would also have to be involved. IDE can form non-proteolytic complexes with other substrates, e.g. alpha-synuclein [16] or a VZV protein precursor [39], which could be sensed by UPR effectors.

At the steady state, IDE deficiency induced a mild UPR with hallmark activation of ATF6 and eIF2α phosphorylation in the absence of PERK activation (Figure 3I-M). Proteotoxic stress can activate other kinases of the integrated stress response phosphorylating eIF2α; for example, proteasome dysfunction is sensed by protein kinase R (PKR) [40]. As IDE may be involved in cytosolic recycling of amino acids from low molecular weight proteins, its absence might also be sensed by general control non-derepressible-2 (GCN2) kinase [41]. Relative enrichment of pathways related to amino acid metabolism in WT islets from HFD-fed mice (Figure 7) would be consistent with such a scenario.

Beta cell proliferation in the presence of a low-level UPR or upon HFD feeding has been observed by multiple authors [42,43][3], with an important role of ATF6 [44][45]. In line with these reports, we document enhanced proliferation of *Ide^-/-^* islet and beta cells, further amplified by massive proteotoxic or metabolic stress. Enhanced proliferation also characterizes *Ide^-/-^* pancreatic alpha cells [46]; although the exact mechanism remains unclear, Merino and colleagues suggested a link with reduced cilia formation, or a role of IDE interaction with PTEN [47]. As discusssed by Cozar-Castellano and colleagues, hyperplasia of, and excessive hormone output by pancreatic endocrine cells upon conditional or global IDE deletion may indicate an important functional role of IDE in secretory processes that remains to be deciphered [21,46].

Proliferation depended on mTORC1 activity and was inhibited by salubrinal, suggesting a requirement for reversing stress-induced eIF2α phosphorylation. In stress situations, mTORC1 can orchestrate an anabolic rescue program, activating amino acid transport and protein synthesis via ATF4 activation [48]; prolonged activation of this program can result in beta cell apoptosis. Elevated serum levels and islet content of proinsulin, observed in HFD-fed *Ide^-/-^* mice, are a hallmark of beta cell dysfunction and suggest unabated mTORC1 activation. Taken together these findings suggest that stressed *Ide^-/-^* beta cells may upregulate an mTORC1-dependent rescue pathway, enhancing proliferation at low but inducing beta cell dysfunction at high metabolic stress levels.

Potentially the most interesting result was the strong upregulation of REG2 mRNA and protein exclusively in infiltrated *Ide^-/-^* islets from NOD mice. REG2 is secreted by beta cells and may act in an autocrine fashion [29]. Treatment of NOD mice with REG2 may reduce the incidence of autoimmune diabetes and increase the beta cell area [32]. In tunicamycin-treated MIN6 insulinoma cells, over-expression of REG2 attenuated Ire1α and PERK in response to tunicamycin while upregulating BIP and at the same time phosphorylation of mTORC1 and of its substrate S6K1 [33]. Thus, the literature links REG2 to an attenuated UPR, reduced inflammation and mTORC1 activation. Accordingly, in our hands, REG2 upregulation is associated with reduced levels of inflammatory cytokines in young NOD mice and protection from autoimmune diabetes, and at steady state with evidence of mTORC1 activation.

However, REG2 is not required for proliferation and in fact downregulated in the presence of massive ER stress. Attenuated inflammation, an ameliorated UPR and a mTORC1-driven regeneration/proliferation program may all underlie protection from autoimmune diabetes. We are aware that proteolytic processing of insulin, the key autoantigen in the NOD model of autoimmune diabetes [49], by IDE may affect disease and have studied T cell responses to insulin in *Ide^-/-^* mice; these studies indicate that altered presentation of insulin peptides by beta cells may also contribute to protection from disease (manuscript in preparation). In any case, beta cell regeneration and an attenuated UPR have both and independently been shown to protect NOD mice from autoimmune diabetes [4,44], rendering a contribution of the present findings in *Ide^-/^*^-^ islet cells to protection plausible.

How could the protective effect of IDE deficiency in NOD mice link to the genetic association of *IDE* with type 2 diabetes? Efforts to develop inhibitors for therapeutic use in this condition [22,50] also assume a detrimental role of IDE, although based on direct inhibition of hormone degradation which may not be the main function of IDE as discussed above. If we assume instead that IDE plays an important but ill-defined role in protein homeostasis and regulation of secretion in endocrine cells, many scenarios protecting from both type 1 and type 2 diabetes can be envisaged. For example, an efficient moderately activated UPR inducing proliferation will be beneficial in both conditions.

The beneficial effect of IDE deficiency on beta cell proliferation and autoimmune inflammation raises the question of the interest of therapeutic Ide inhibition. IDE inhibitors have previously been proposed to be of interest to ameliorate glucose metabolism in type 2 diabetes patients, and more recently also to reduce autoimmune inflammation in the NOD model. Maianti et al found that an inhibitor competing with substrate binding ameliorated glucose tolerance in an OGTT [22], while we found that administration of an active site inhibitor compromised glucose tolerance in the same test [23]. The reason for this discrepancy remains unclear. A novel inhibitor designed by Nash and colleagues binding to the central substrate binding cavity reportedly provides protection from diabetes in NOD mice, seemingly consistent with our data [50]; REG2 expression was not studied by these authors.

However, in contrast to our results, this inhibitor improves rather than compromises glucose tolerance in HFD-mice and does not induce weight gain, therefore probably does not result in mTORC1 activation. Collectively these findings indicate that our mechanistic understanding of the effects of IDE inhibition remains poor. Our finding that IDE deficiency activates not only proliferation but also an anti-inflammatory pathway (Figure 8) indicates that biological effects of IDE unrelated to insulin degradation should be of significant interest for future research.

## 4. MATERIALS AND METHODS

### 4.1. Mouse models

*Ide^-/-^* mice on a C57BL/6 background were obtained from S. Guenette [8]. NOD.*ide* congenic stock was generated using a marker-assisted breeding approach to speed up the introgression of the *Ide*-disrupted gene from the C57BL/6 congenic mice onto a NOD genetic background [37]. Specifically, a panel of 36 informative microsatellite markers was chosen to cover all the insulin-dependent diabetes (Idd) susceptibility loci of the NOD mice known at the time (I*dd1* to *Idd20*). Although microsatellites analyses confirmed that all NOD *Idd* loci had been fixed after 5 generations, another 5 back-crosses to NOD mice were performed. The presence of the desired *Ide^+/+^* or *Ide^-/-^* genotypes was routinely verified by PCR using the primers, WT Forward: 5’-ATCTG TGTCA GGAGG AGGGA C, WT Reverse: 5’-CAGGG TAGGG AAGTC AAGGT TAC, and NEO Forward: 5’-GGGCG CCCGG TTCTT TDTTTGT C, NEO Reverse: 5’: TTGGT GGTCG AATGG GCAGG T. C57BL/6J, C57BL/6J *Ide^-/-^*, and NOD *Ide^+/+^*, NOD *Ide^-/-^*, and NOD *Ide^-/-^ Rag2^-/-^* were bred and housed in an ambient temperature room with 12/12 h light/dark cycles and specific pathogen-free conditions. Ethical approval for animal experimentation in this project was accorded by the Ministère de l’enseignement, de la recherche et de l’innovation on October 8, 2020, for a duration of 5 years under the number APAFIS#27441-2019080809515121.

### 4.2 Insulitis scoring

Pancreata were fixed in 4% formalin for at least 2 h, embedded in paraffin overnight and mounted in a paraffin block. Four µm paraffin sections were mounted onto Superfrost™ slides coated with albumin, dried, deparaffinized and re-hydrated in 100%, 90%, 80% alcohol baths. Sections were stained in hematoxylin and eosin, each for 2 min, and mounted with EUKIT and a coverslip. For each pancreas, 30 islets were scored for insulitis using the following classification [38]: 0: normal, 1: peri-insulitis, 2: moderate insulitis (less than 50% of mononuclear cells) 3: severe insulitis (more than 50% of mononuclear cells and/or loss of islet architecture).

### 4.3. Spontaneous Diabetes Incidence

The onset of diabetes was defined as two positive urine glucose tests, confirmed by a glycemia >200 mg/dL. Glucose tests and measure of glycemia were performed in a blind fashion.

### 4.4. Oral glucose tolerance test (OGTT)

OGTT experiments were performed on *Ide^+/+^* and *Ide^-/-^* NOD mice as described in reference [23]. Following a 6 h fasting period, basal blood glucose was determined and a bolus of glucose at 2 g/kg was administered by gavage. Blood glucose levels were determined using an Accu-Check glucometer (Roche). Plasma concentrations were measured by ELISA kit (10-1247-01, Mercodia) in blood samples collected 15 min before and 15, 90 and 180 min after glucose administration.

### 4.5. Estimation of islet volume

Islets from 15 mice aged 8 weeks per group were stained for insulin and imaged by fluorescence microscopy. An average of 40 plans per islet were acquired. Islet volume was determined using Icy software and the plug-in ROI Statistics developed by members of the Biological Image Analysis unit of the Institut Pasteur, Paris.

### 4.6. Pancreatic islet preparation

Ice-cold collagenase P (Roche; 0.76 mg/ml) was injected into the pancreatobiliary duct. The instilled pancreas was digested for 10 min at 37°C, digestion was stopped by addition of cold HBSS with 10% FCS. The islets were handpicked under the microscope.

### 4.7. IAPP quantification in serum

To determine IAPP levels in blood, undiluted serum from female mice of different age was analyzed using a commercial ELISA kit (TSZ Biological, North Brunswick, NJ, USA).

### 4.8. Glucose-stimulated insulin secretion (GSIS)

15 similar size islets were preincubated in fresh DMEM with 10% FCS without glucose for 90 min, followed by 1 h of incubation in DMEM with 2.8-or 28-mM glucose. Insulin content in supernatants was quantified using an insulin ELISA (10-1247-01, Mercodia).

### 4.9. RNA preparation and qPCR analysis

Total RNA of isolated islets was extracted using NucleoSpin RNA XS kit (740902.50, Macherey-Nagel), then normalized RNA was transformed into cDNA using the High-capacity cDNA reverse transcription kit (43-688-14, Applied Biosystems). QPCR was performed on a 7900HT fast real-time PCR system (Applied Biosystems) to identify gene expression.

Relative gene expression levels were expressed as ΔΔCt (fold change), and the primers used are listed in Supplementary Table 3.

### 4.10. Digestions with recombinant IDE

Recombinant mouse REG2 (2098-RG, R&D Systems) and human insulin (11376497, Roche) were digested with recombinant WT or protease-dead (cf-E111Q) human IDE enzyme dead enzyme (24) in 20 mM Hepes pH 7.2, 50 mM NaCl buffer at a ratio IDE: INS ratio (w/w) of 1:1, or a ratio IDE: REG2 of 1:10. Enzymes and substrates were incubated with end-over-end rotation at 37°C for 5 s to 5 min for insulin, and for 30 min to 8 h for REG2. Samples were resolved by 4-12% NuPAGE Bis-Tris gels (NP0321BOX, Invitrogen), stained with SYPRO® Ruby Protein Gel Stain (S12000, Molecular Probes) and visualized using a UV source.

### 4.11. Proteomic analysis of NOD islets

Samples from 3 *Ide^+/+^* and 3 *Ide^-/-^* mice aged 10 weeks (∼100 islets per sample) were prepared, digested and then resuspended in 10% ACN, 0.1% TFA in HPLC-grade water. Each sample was injected three times in a nanoRSLC-Q Exactive PLUS (RSLC Ultimate 3000) (Thermo Scientific). Peptides were loaded onto a µ-precolumn (Acclaim PepMap 100 C18, cartridge, 300 µm i.d.×5 mm, 5 µm) (Thermo Scientific), and separated on a 50 cm reversed-phase liquid chromatographic column (0.075 mm ID, Acclaim PepMap 100, C18, 2 µm) (Thermo Scientific). Peptides were eluted and analyzed by data dependent MS/MS, using top-10 acquisition method. Peptides were fragmented using higher-energy collisional dissociation (HCD). The MS files were processed with the MaxQuant software version 1.6.17 and searched with the Andromeda search engine against the UniProtKB/Swiss-Prot Mus Musculus database (release of February 2021, 17071 entries). Proteins were quantified according to the MaxQuant label-free algorithm using LFQ intensities. Statistical and bioinformatic analysis were performed with Perseus software (version 1.6.14) (www.perseus-framework.org). We performed a two-sample t-test with a p-value threshold of 0.05. The differentially expressed proteins were subjected to bioinformatic analysis using EnrichR software (https://maayanlab.cloud/Enrichr/) for enrichment of GO terms using GO Biological Process library from 2021 (Supplementary Table 2). TOP10 enriched terms were ranked using adjusted p-value ranking. Volcano-plot and enrichment bar-plot were made using Python (version 3.8.3).

### 4.12. ER stress

Mice either received an intraperitoneal injection of PBS (10 mL/kg, control group) or tunicamycin (T7765, Sigma-Aldrich; 2 mg/kg) or two injections of rapamycin (R-5000, LC laboratories; 5 mg/kg at day1 and day 2) or tunicamycin + rapamycin, or tunicamycin + salubrinal (SML0951, Sigma-Aldrich; 1.5 mg/kg). Mice were sacrificed 24 h to 48 h post-injection.

### 4.13. Immunoblots

Isolated islets were homogenized in RIPA lysis buffer (89900, Thermo scientific) containing protease (11836170001, Roche) and phosphatase inhibitor cocktails (4906845001, Roche). Samples were resolved by 4-12% NuPAGE Bis-Tris gels (NP0321BOX, Invitrogen), electrotransferred to polyvinylidene difluoride membranes using I-Blot2 (IB21001, Invitrogen), and probed overnight at 4°C with primary antibodies against phospho-PERK, total PERK, phospho-eIF2α, total eIF2α, ATF6, PCNA and β-actin. Detection was performed using HRP-labelled anti-rabbit/mouse IgG and developed with SuperSignal West Femto maximum sensitivity substrate (PI34095, Thermo scientific) or West Pico plus chemiluminescent substrate (34580, Thermo scientific), then quantified using ImageJ Software. Antibodies are listed in Supplementary Table 4.

### 4.14. HFD treatment and follow up

Seven-week-old female *Ide^+/+^* and *Ide^-/-^* C57BL/6J mice were fed a standard chow (VRF1, Special Diets Services) or HFD (D12492, Brogaarden) for 7 weeks. Body weight was recorded once a week, and non-fasting glucose concentrations were monitored weekly using Accu-Check glucometer (Roche). Plasma and islet insulin and proinsulin concentrations were measured by commercial ELISA kits (10-1247-01 and 10-1232-01, respectively; Mercodia). For RNAseq analysis of handpicked islets, mice were fed a HFD for 2 weeks.

### 4.15. Immunocytology

Handpicked islets were seeded on SuperFrost Gold Plus microscope slides. For Ki67 staining, islets were fixed with 4% PFA, permeabilized with 0.1% Triton and stained with anti-Ki67, anti-insulin was used to identify β-cells. Nuclei were stained with DAPI (D1306, Thermo Fisher Scientific). Image acquisition was performed using a Leica SP8 confocal microscope. Ki67 positive cells and islet volumes were estimated with Icy Software.

### 4.16. EdU staining

EdU was intraperitoneal injected (100 µg/g) into mice. The pancreas was harvested 1 h later, fixed in Antigenfix (P0014, Diapath) and embedded in paraffin blocks. Sections (5-μm-thick) were cut, deparaffinized in xylene, rehydrated followed by antigen retrieval (208572, Abcam). EdU staining was conducted using the Click-iT^TM^ EdU imaging kit (C10337, Invitrogen) according to the manufacturer’s protocol. Anti-insulin was used to identify β-cells. Slides were mounted with Vectashield mounting media (NC9532821, Vector Laboratories Inc). For each mouse eight randomly selected sections were included for analysis. Images were captured using a Leica SP8 SMD confocal microscope and EdU-positive β-cells were counted.

### 4.17. RNAseq analysis

Total RNA samples were sent for library preparation and sequencing on an Illumina Nextseq 500 instrument. A primary analysis based on AOZAN software (ENS, Paris) was applied to demultiplex and control the quality of the raw data (based of FastQC modules/version 0.11.5). Fastq files were aligned using STAR algorithm (version 2.7.6a), on the Ensembl Mus musculus GRCm38 reference, release 101. Reads were then counted using RSEM (v1.3.1) and the statistical analyses on the read counts were performed with R (version 3.6.3) and the DESeq2 package (version 1.26.0) to determine the proportion of differentially expressed genes (DEGs) between two conditions. Cluster analysis was performed by hierarchical clustering using the Spearman correlation similarity measure and ward linkage algorithm. Heatmaps were made with the ggplot2 (version 3.3.5) and NMF (version 0.23.0) packages and a custom color palette from the RColorBrewer package (version 1.1-2). DEGs with a minimal overlap of 3, a P-value cutoff of 0.01 and a minimal enrichment of 1.5 were selected for the Pathway & Process Enrichment analysis. Input genes lists were included: GO Molecular Functions; GO Biological Processes, Reactome Gene Sets, KEGG Pathway; GO Cellular Components. The output lists of terms were then filtered to select pathways with adjusted p-values (q-values) <0.01. The criteria to select pathways to be plotted are described in figure labels.

### 4.18. Data availability

The mass spectrometry proteomics data have been deposited to the ProteomeXchange Consortium via the PRIDE partner repository (49) with the dataset identifier PXD034826 and 10.6019/PXD034826. The GEO accession number for RNAseq datasets is GSE207797. The data used to generate the results in this paper are available as source data. All data and mouse lines included in this study are available from the corresponding author upon reasonable request.

### 4.19. Statistical analysis

Statistical analyses were carried out using GraphPad Prism 8. Statistical differences were evaluated using two-tailed unpaired Student t test for comparisons of one variable between two groups, one-way ANOVA and appropriate post hoc analyses for comparisons of one parameter between multiple groups and two-way ANOVA with post hoc statistical adjustment for comparisons of two or more parameters between multiple groups. A p value of less than 0.05 (*P < 0.05, **P < 0.01, and ***P <0.001) was considered statistically significant.

## Author contributions

SZ, EWE: investigation, visualization; AM, MAB, JL, KR, AY, BB: investigation; BSP, CIG, FXM: formal analysis; JD, SF: conceptualization; PVE: conceptualization, funding acquisition, administration, supervision, visualization, writing, editing. The position of the first listed author was determined by flipping a coin.

## Acknowledgements

This work was supported by a grant from the European Foundation for the Study of Diabetes EFSD/Lilly Program 2017), ANR-18-CE92-0008-01 from the *Agence Nationale de Recherche*, and grant EQU201903007853 from the *Fondation pour la Recherche Médicale*. We are greatful to Vinson Liang and Wei-Jen Tang, University of Chicago, who provided recombinant IDE enzyme, with support through grant NIH R01 GM121964.

---

**Figure S1 related to Figure 1. Ide^-/-^islets are devoid of staining for IDE in fluorescent microscopy.** Dissociated islet cells (**A**) or hand-picked islets (**B**) from WT and *Ide^-/-^* C57BL/6 mice were stained with a mouse antibody (A) or goat antibodies (B) against IDE and Alexa594-coupled secondary goat (A) or donkey (B) secondary antibodies and analyzed by fluorescence microscopy.

**Figure S2 related to Figure 2. REG2 induction in *Ide^-/-^* islets is due to inflammation and not to absence of its digestion by IDE. (A, B)** expression of genes related to the UPR and to proliferation in islets pre-incubated or not overnight was evaluated as in Figure 2. **(C, D)** Recombinant mouse REG2 and human insulin were digested for the indicated duration with recombinant WT or protease-dead human IDE. Active IDE rapidly digests insulin but is completely unable to degrade REG2.

**Figure S3 related to Figure 2. Effect of proteotoxic stress on UPR, proliferation and regeneration markers.** *Ide^-/-^* and *Ide^+/+^* 12-week-old C57BL/6 **(A-C;** N = 3) or 10-week-old NOD **(D-I;** N = 4) mice were treated for 2 days with tunicamycin alone or combined with rapamycin before collection of islets and analysis for gene expression by RT-qPCR. Results are expressed as fold induction comparing the two conditions indicated in the panels. Data are mean values ± SEM and were analyzed by ANOVA test for statistical significance. *p < 0.05, **p < 0.01, ***p < 0.001, ****p < 0.0001, by ANOVA test.

**Figure S4 related to Figure 4. Effect of IDE deficiency on inflammatory gene expression in female and male NOD mice. (A)** mRNA expression of IFIT3, MCP-1, NLRP-3, IF27L2A, and IFI47 was assessed by RT-qPCR in the islets of *Ide^-/-^* and *Ide^+/+^* NOD mice at 4, 6, 9, 12, and 15 weeks of age. Results are expressed as ratio of expression between *Ide^-/-^*and *Ide^+/+^* mice. Data are mean values ± SEM of 8 independent mice for each group. **p < 0.02, and ***p < 0.01 by Student’s t test-test. **(B)** experiment as in **(A)** but examining expression of the genes indicated in islets from 12-week-old male NOD mice, N = 5.

**Figure S5 related to Figure 5. Effect of acute proteotoxic stress on (pro)insulin production and secretion.** 11-week-old female *Ide^-/-^* and *Ide^+/+^* NOD mice were treated by tunicamycin alone or combined with rapamycin for 2 days before measuring serum concentrations **(panels A, B)** and islet content **(panels C, D)** of insulin **(A, C)** and proinsulin **(B, D)**. Data are mean values ± SEM from 2 experiments each with 4 mice per group. *p < 0.05, **p < 0.01, ***p < 0.001, ****p < 0.0001), by Student’s t-test.

**Figure S6 related to Figure 7.** Expression of selected genes in *Ide^+/+^* and *Ide^-/^*^-^ islets of mice subjected to metabolic stress. **(A, B)** Color heatmap (Z-score) of key genes selected from the list of differentially expressed genes (adjusted p-value<0.01, l2fc>2) and classified into general cellular processes: cell cycle **(A)** and immune system **(B)**.

