## Supplementary figures and images for "Pancreatic islet cell stress induced by insulin-degrading enzyme deficiency promotes islet regeneration and protection from autoimmune diabetes"

### Supplemental Figure 1

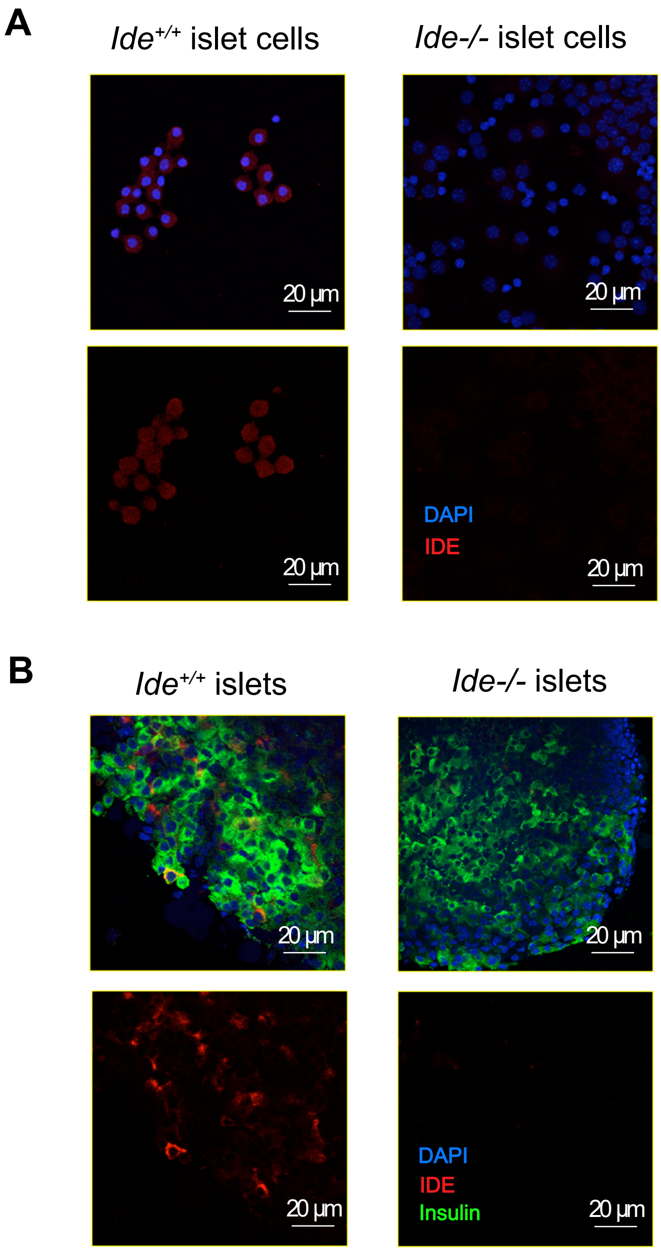

### Supplemental Figure 2

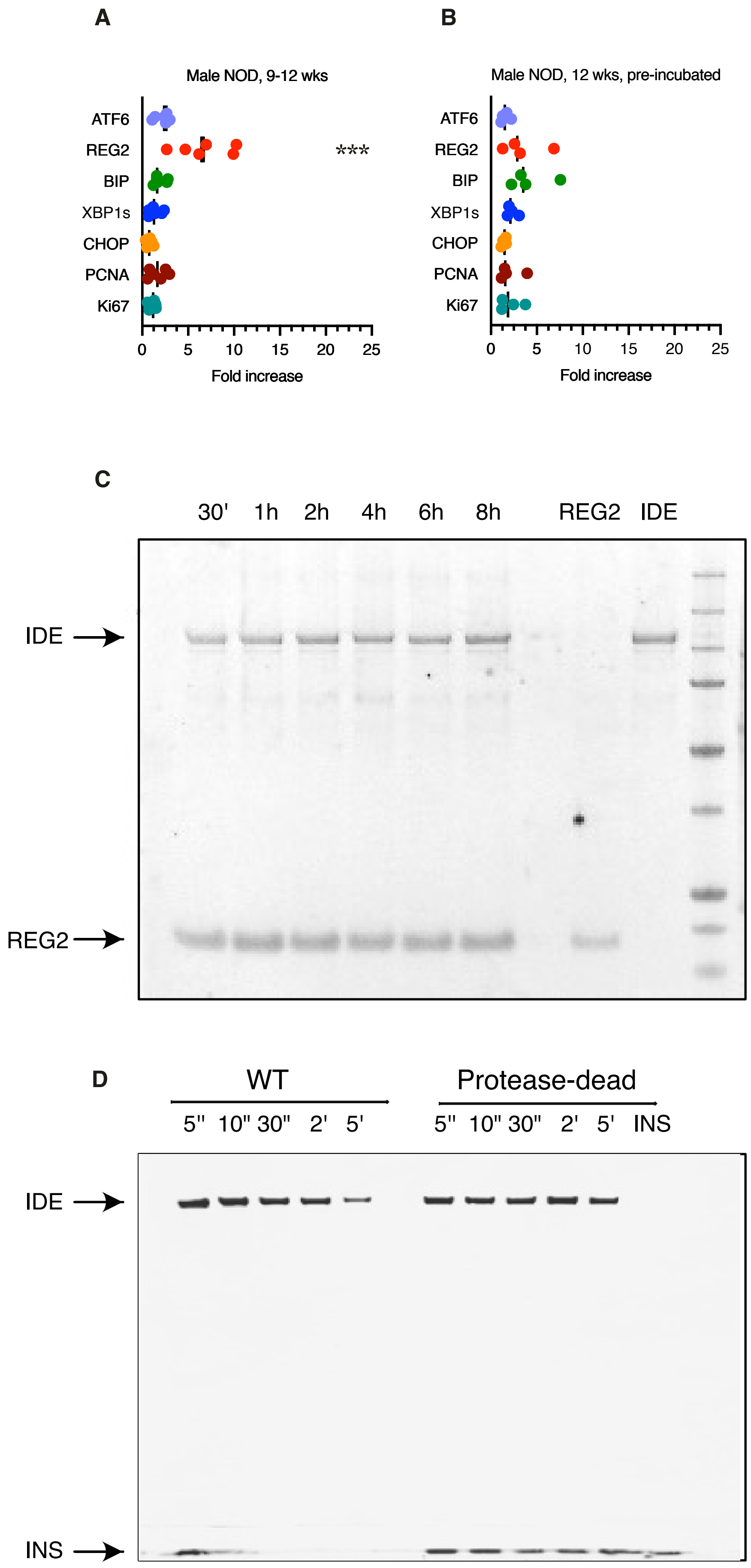

### Supplemental Figure 3

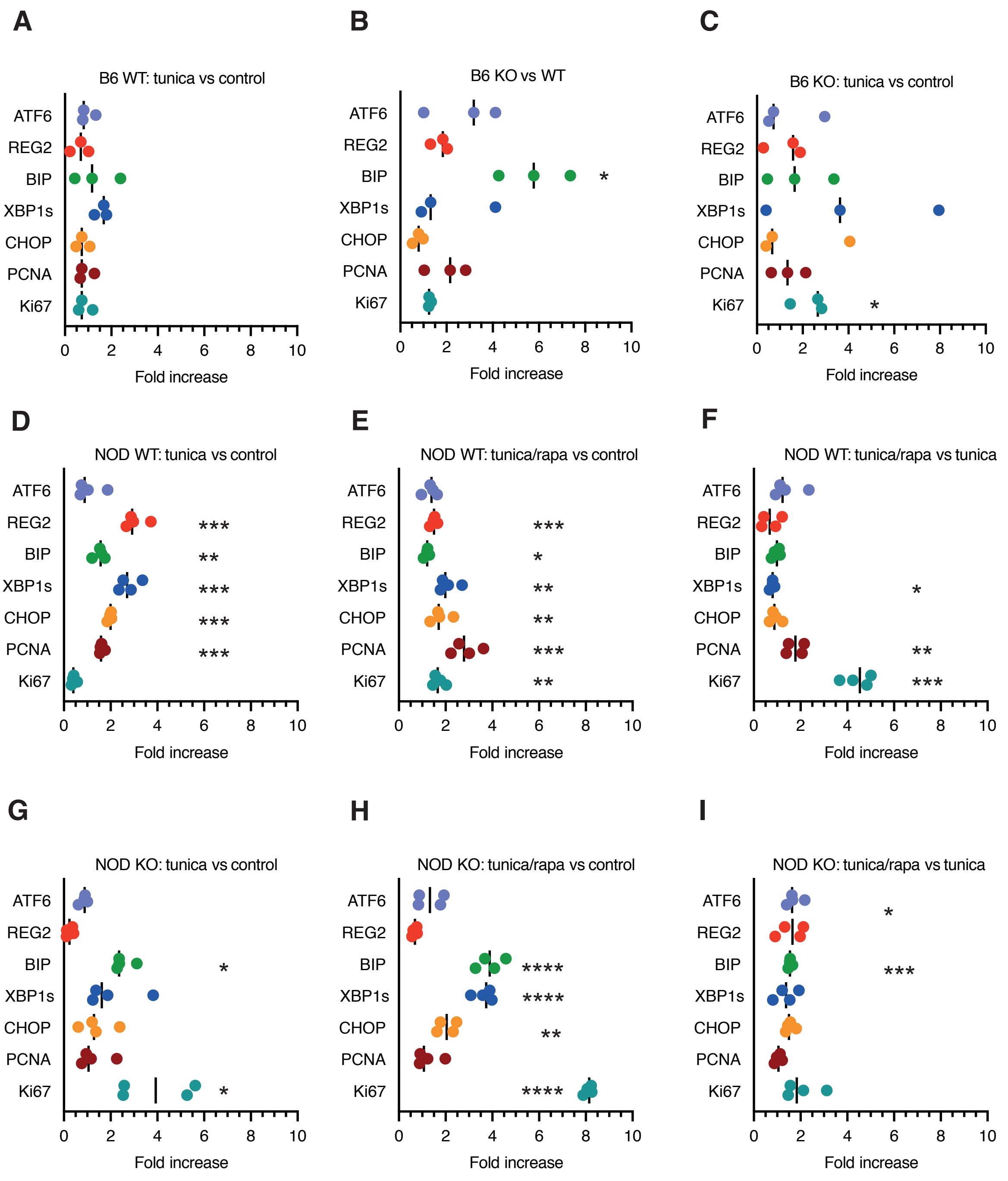

### Supplemental Figure 4

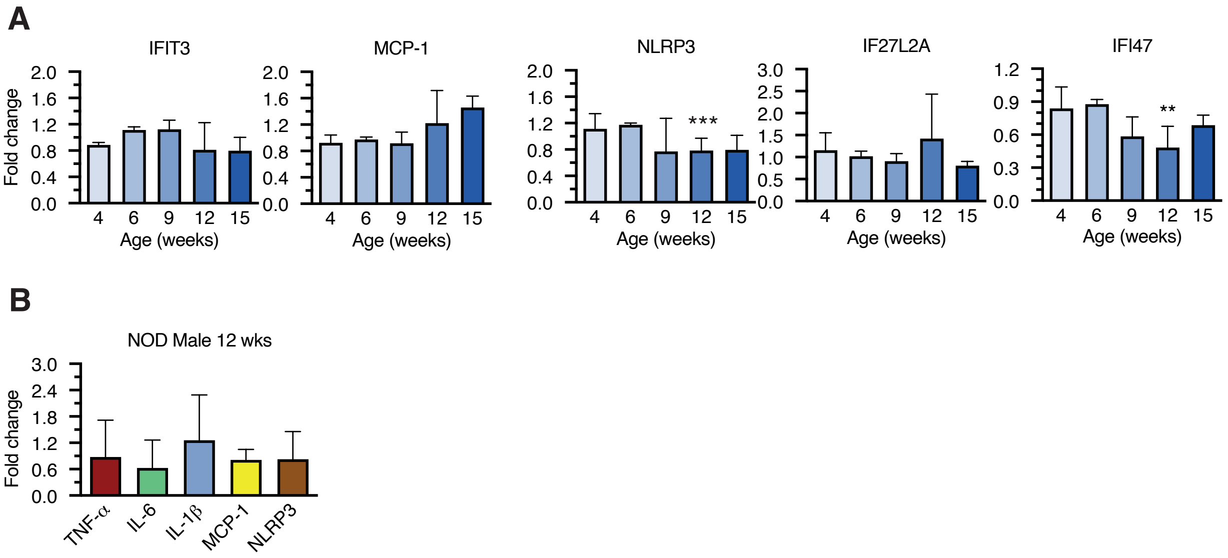

### Supplemental Figure 5

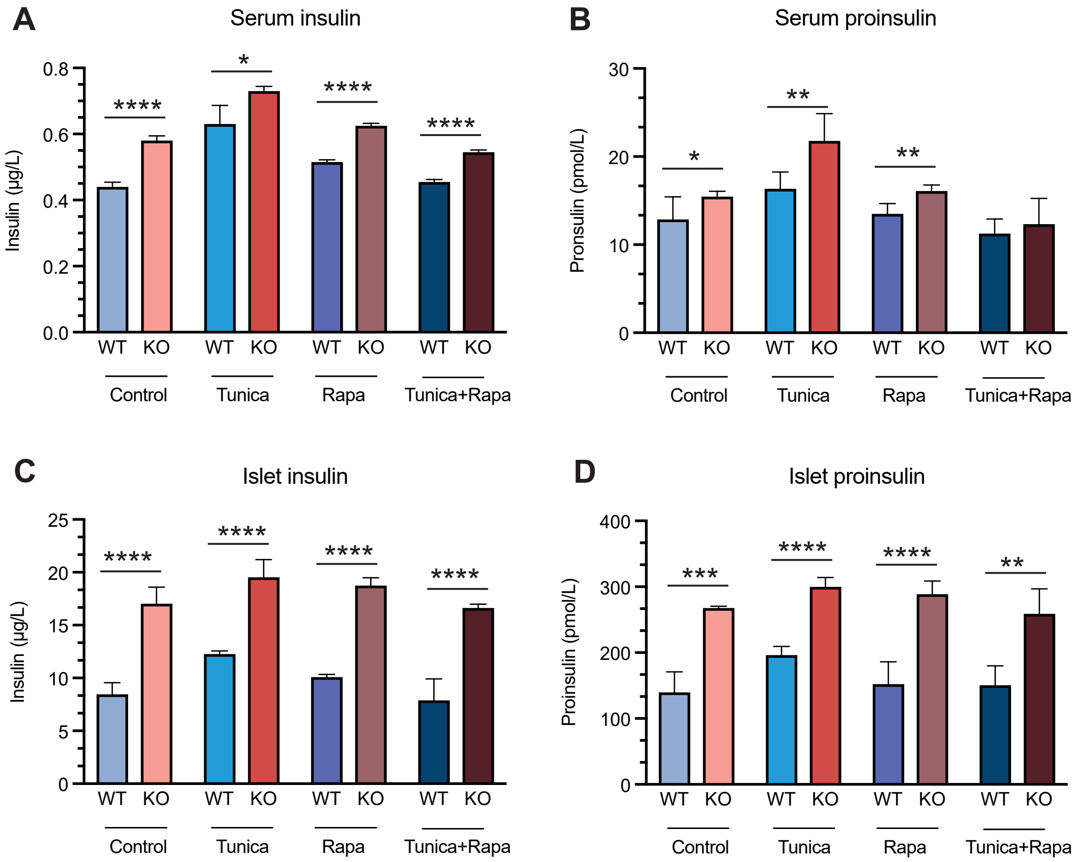

### Supplemental Figure 6

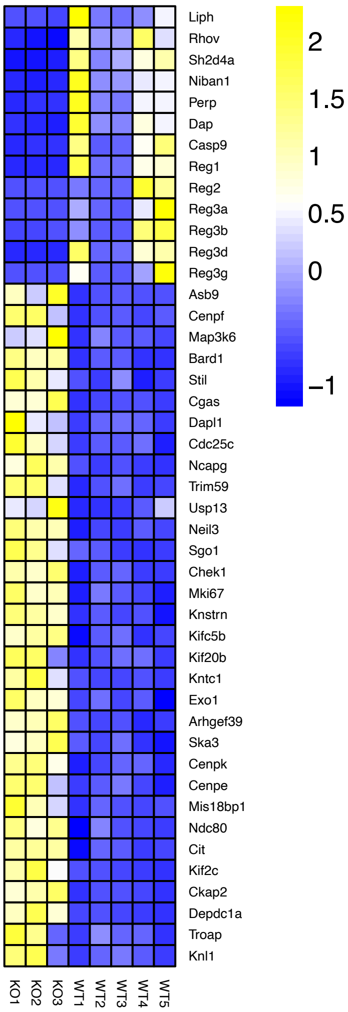

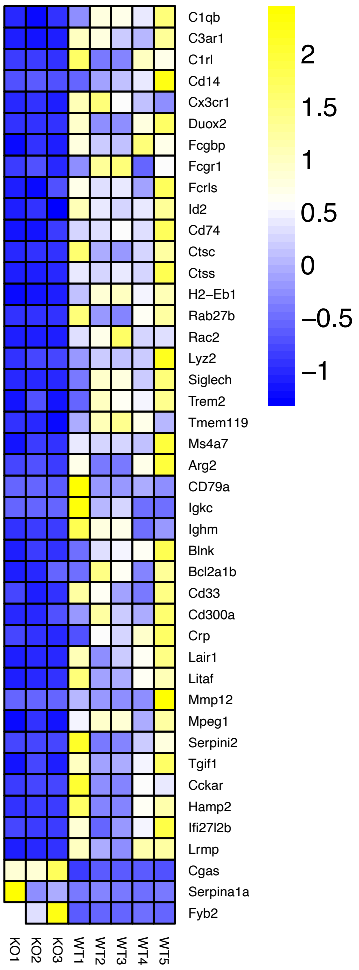
