## Supplemental Table 3 for "Pancreatic islet cell stress induced by insulin-degrading enzyme deficiency promotes islet regeneration and protection from autoimmune diabetes"

Supplementary Table 3.

Genes Primers	5’ to 3’

Ki67 forward	GGCGGCCAGAGCTAACTT

Ki67 reverse	TGACACTACAGGCAGCTGGA

PCNA forward	GAACCTCACCAGCATGTCCA

PCNA reverse	AATTCACCCGACGGCATCTT

BiP forward	GTGTGTGAGACCAGAACCGT

BiP reverse	GTTCTTGAACACACCGACGC

REG2 forward	TGCCAGAACATGAATGCA

REG2 reverse	TCAGCGATACACAATAACCAC

CHOP forward	CCGGAACCTGAGGAGAGAGTG

CHOP reverse	TCTTCGTTTCCTGGGGATGAG

XBP1s forward	CTGAGTCCGCAGCAGGTG

XBP1s reverse	GACCTCTGGGAGTTCCTCCA

ATF6 forward	AGCATACAGCCTGCGCCCAC

ATF6 reverse	CACCGGGCTGCTGCTCACCACAG

MCP-1 forward	AAAACACGGGACGAGAAACCC

MCP-1 reverse	ACGGGAACCTTTATTAACCCCT

IL-6 forward	TTCCATCCAGTTGCCTTCTTG

IL-6 reverse	GGGAGTGGTATCCTCTGTGAAGTC

NLRP3 forward	ATTACCCGCCCGAGAAAGG

NLRP3reverse	TCGCAGCAAAGATCCACACAG

TNF-α forward	GGCAGGTTCTGTCCCTTTCAC

TNF-α reverse	TTCTGTGCTCATGGTGTCTTTTCT

IL-1β forward	GCAACTGTTCCTGAACTCAACT

IL-1β reverse	ATCTTTTGGGGTCCGTCAACT

OAS3 forward	TCTGGGGTCGCTAAACATCAC

OAS3 reverse	GATGACGAGTTCGACATCGG

IFI47 forward	TCTCCAGAAACCCTCACTGGT

IFI47reverse	TCAGCGGATTCATCTGCTTCG

IFIT3 forward	CCTACATAAAGCACCTAGATGGC

IFIT3 reverse	ATGTGATAGTAGATCCAGGCGT

IFI27L2A forward	TCAGCAGGGGTCCTTGGACTCTC

IFI27L2A reverse	CATCTCCTGCGTAGTCTGTACAGGC

GAPDH forward	CCGTAGACAAAATGGTGAAGG

GAPDH reverse	CGTGAGTGGAGTCATACTGGA
