## Supplemental Table 4 for "Pancreatic islet cell stress induced by insulin-degrading enzyme deficiency promotes islet regeneration and protection from autoimmune diabetes"

Supplementary Table 4.

Antibodies	Source	Dilution

Rabbit monoclonal anti-phosphor-PERK	Cell signaling Cat# 3179	1:1000

Rabbit monoclonal anti-PERK	Cell signaling Cat# 3192	1:1000

Rabbit polyclonal anti-phosphor-eIF2α	Cell signaling Cat# 9721	1:1000

Rabbit polyclonal anti-eIF2α	Cell signaling Cat# 9722	1:1000

Mouse monoclonal anti-ATF6	Novus Biologicals Cat# NBP1-40256	1:500

Rabbit monoclonal anti-PCNA	Abcam Cat# ab92552	1:1000

Mouse monoclonal anti-β-actin	Santa Cruz Cat# sc-47778	1:1000

Alexa Fluor® 647 Mouse anti-Ki-67	BD Biosciences Cat# BD558615	1:10

Guinea pig polyclonal anti-insulin	Arigo biolaboratories Cat# ARG62502	1:50

Rabbit polyclonal anti-Amylin Santa Cruz Cat# sc-20936	1:1000

Mouse anti-IDE	Santa Cruz Cat# sc-27266	1:100

Goat anti-IDE	Santa Cruz Cat# sc-393887	1:100

HRP-labelled anti-rabbit IgG	Cell signaling Cat# 7074	1:3000

HRP-labelled anti-mouse IgG	Cell signaling Cat# 7076	1:3000

Goat anti-guinea pig Alexa Fluor® 594 Invitrogen Cat# A-11076	1:200

Goat anti-guinea pig Alexa Fluor® 488	Invitrogen Cat# A-11073	1:200

Donkey anti-goat Alexa Fluor® 594   Invitrogen Cat#A-11058	1:400

Goat anti-mouse Alexa Fluor® 594	Invitrogen Cat#A-21202	1:400
